# Surface electrostatics govern the emulsion stability of biomolecular condensates

**DOI:** 10.1101/2020.04.20.047910

**Authors:** Timothy J. Welsh, Georg Krainer, Jorge R. Espinosa, Jerelle A. Joseph, Akshay Sridhar, Marcus Jahnel, William E. Arter, Kadi L. Saar, Simon Alberti, Rosana Collepardo-Guevara, Tuomas P.J. Knowles

## Abstract

Liquid–liquid phase separation underlies the formation of biological condensates. Physically, such systems are microemulsions which have a general propensity to fuse and coalesce; however, many condensates persist as independent droplets inside cells. This stability is crucial for their functioning, but the physicochemical mechanisms that control the emulsion stability of condensates remain poorly understood. Here, by combining single-condensate zeta potential measurements, optical microscopy, tweezer experiments, and multiscale molecular modelling, we investigate how the forces that sustain condensates impact their stability against fusion. By comparing PR_25_:PolyU and FUS condensates, we show that a higher condensate surface charge correlates with a lower fusion propensity, and that this behavior can be inferred from their zeta potentials. We reveal that overall stabilization against fusion stems from a combination of repulsive forces between condensates and the effects that surface electrostatics have on lowering surface tension, thus shedding light on the molecular determinants of condensate coalescence.

## Introduction

Solutions of multivalent macromolecules, including multidomain proteins and those comprising intrinsically disordered regions, peptides, and nucleic acids, have the ability to undergo demixing through liquid–liquid phase separation (LLPS) (*1*, *2*). LLPS enables the formation of condensed-liquid droplets, which coexist with a dilute aqueous phase (*3–5*). This process occurs when the energetic gain that a biomolecular system incurs by forming a densely connected condensed liquid network surpasses the entropic cost of demixing and reducing its available number of microstates. In living cells, LLPS has been shown to underlie the formation of biomolecular condensates which function as membraneless organelles. This process provides a mechanism for the spatiotemporal control (*6*) of several vital processes (*7*), including RNA processing and stress signaling (*8*, *9*). Moreover, aberrant LLPS, often involving liquid-to-solid transitions, has been implicated in the emergence of various neuropathologies, age-related diseases, and cancer (*10–12*).

Biomolecular condensates are highly diverse systems, both in terms of composition, size, and behavior. Not only is the range of different proteins and nucleic acids that can undergo LLPS both *in vitro* and in cells vast (*8*, *10*, *13*), but mounting evidence also suggests that the detailed chemical nature of the interactions that drive these diverse systems to phase separate spans a range of charge–charge, cation–π, π–π, polar, hydrophobic, and even hybrid interactions (*2*, *14*, *15*). A unifying feature of intracellular LLPS is that, in most if not all cases, condensate formation is driven by a combination of both electrostatic and non-ionic interactions; however, the exact balance amongst these forces is diverse and dictated not only by the chemical makeup of the biomolecules in question but also by the microenvironment they are exposed to (*15*).

A similar richness in the behavior of biomolecular condensates is exemplified by the significant variation in their fusion and coalescence propensities, with wide-ranging functional implications (*16–18*). For instance, processes like stress adaptation and signaling (*7*, *19*) depend on the ability of phase-separated liquid drops to remain stable against fusion for varying periods of time that range from seconds to hours. In other cases, for example in the nucleoli, fusion of multiple droplets into a single large condensate phase may be critical for clustering of RNA and subsequent functionality (*20*). Previous work has suggested that active chemical and biological processes in cells may operate to prevent droplet coalescence (*21*). Observations of protein liquid condensates coexisting *in vitro* without undergoing fusion also suggest, however, that passive mechanisms exist that prevent condensates from rapidly fusing and clustering (*22*, *23*). These examples highlight the relevance of investigating the molecular mechanisms that control stability of condensates against fusion from a fundamental perspective. Of particular interest in this respect is the question of how the diverse molecular forces that sustain condensate LLPS impact their propensity to coalesce or to remain stable against fusion.

Physically, biomolecular condensates are water-in-water emulsions with a low surface tension, similar to other polyelectrolyte coacervate systems or colloidal assemblies (*21*). A specific quantity of interest that has long been used to describe the stability of such emulsions against coalescence, coagulation, and clustering is the zeta potential (*24–26*)— the electro-kinetic potential at the edge of the interfacial double layer coating the surface of any charged particle (Figure 1A). In particular, low absolute zeta potentials, *i.e.*, usually smaller than 30 or 40 mV in absolute value, tend to be associated with emulsions that fuse (*27*, *28*). Outside this regime, electrostatic repulsion is suggested to enable emulsions to remain stable against fusion (*29*, *30*). Based on these observations, we sought to examine if the zeta potential of protein condensates could be established as a new parameter to assess and predict the propensity of condensates to fuse and coalesce, and to infer electrostatic properties of condensate surfaces. Moreover, we aimed to rationalize how mesoscale zeta potential values emerge from the distribution and molecular organization of proteins, water, and ions in and around condensates.

**Figure 1.**
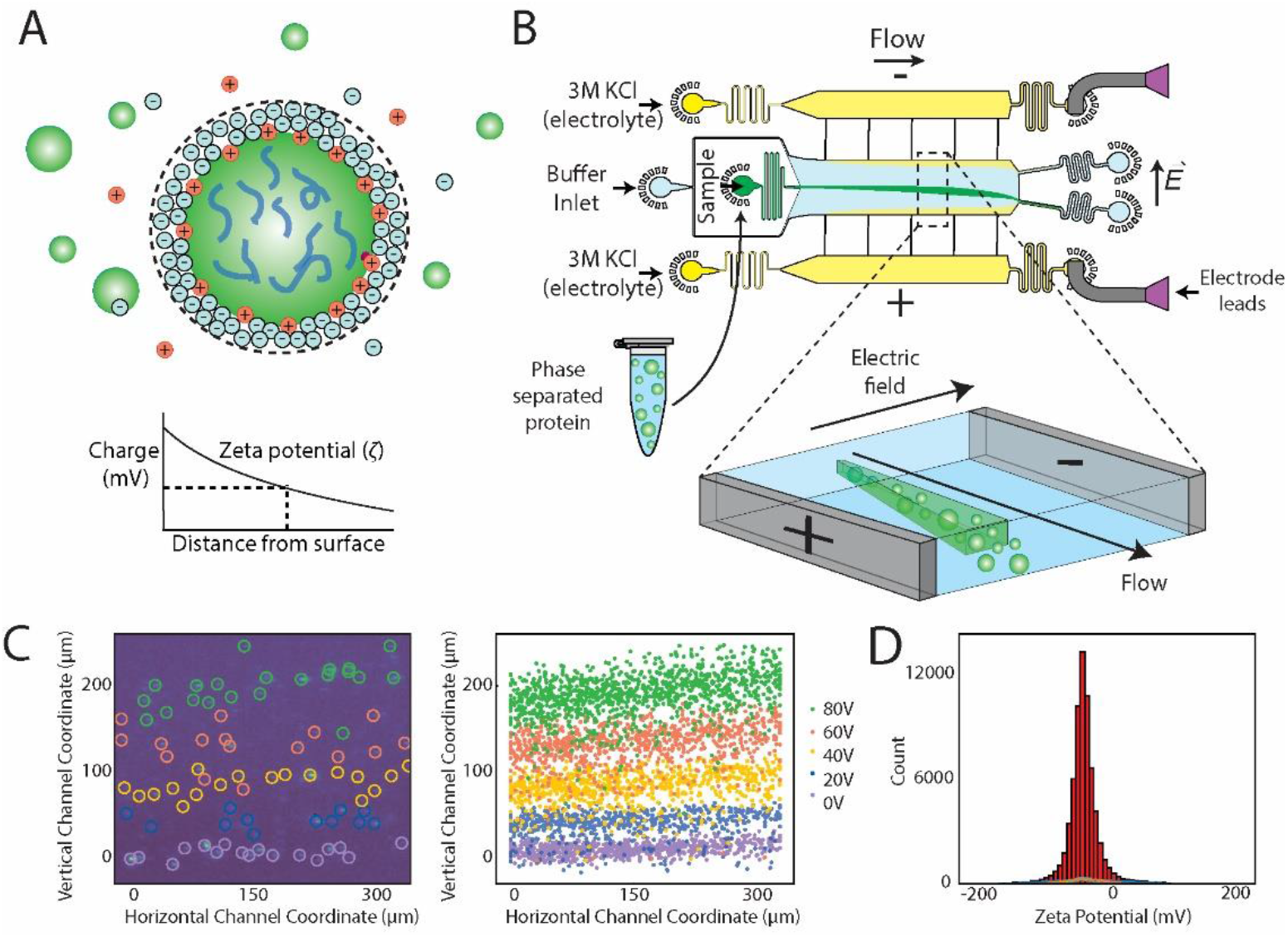
Overview of the microfluidic platform for quantifying single-condensate zeta potentials. **(A)** Schematic of the zeta potential of a protein condensate, which is the electrical potential at the edge of the ion layer surrounding a particle, denoted by the dashed lines. **(B)** Schematics of the μFFE device used to carry out the single-droplet zeta potential measurements. Phase separated droplets were introduced into the 3D free-flow electrophoresis device through a central injection port, preventing any contact between the condensates and the surface of the channel. The condensates were then deflected by applying a constant voltage and positions quantified as a measure of electrophoretic mobility to calculate zeta potentials. **(C)** Left panel: Overlaid images from multiple voltage applications in the range from 0–80 V, depicting individual protein condensates as they move through the image frame. Right panel: Tracked coordinates of detected condensates at each voltage in the range between 0 and 80 V; these coordinates were used to calculate the zeta potential (see Materials and Methods). **(D)** Each individual condensate was analyzed to yield single-droplet zeta potential distributions, represented as the sum of all obtained measurements across all voltages applied. Cityscapes at the bottom of each filled histogram are histograms derived from single voltage measurement.

To this end, we devised a microfluidic approach that enables measurement of zeta potentials at the resolution of individual condensates. We correlated these measurements with the propensity of condensates to fuse and coalesce using epifluorescence and brightfield microscopy, as well as optical tweezer experiments. Subsequently, to obtain a molecular understanding of our experimental observations, including characterizing the behavior of counterions in and out of condensates, we developed a multiscale molecular modelling strategy that equilibrates protein condensates at coarse-grained resolution and then back-maps them to the atomistic level, including explicit solvent and ions. We show that zeta potentials obtained for various biomolecular condensates correlate well with their propensity to fuse, coalesce, and cluster. Our multiscale molecular dynamics simulations help to elucidate the molecular origin of the different zeta potential values—linking fusion propensities to the modulation of the surface tension of condensates via surface electrostatics. These results establish the zeta potential as a fundamental quantity to infer the tendency of biomolecular condensates to fuse and coalesce and rationalize it from the molecular organization of charged species in the system.

## Results

### Single-condensate zeta potential measurements

To quantify the zeta potential of biomolecular condensates experimentally, we developed a single-particle microfluidic approach based on free-flow electrophoresis (μFFE) using a 3D device, that enables in-solution quantification of zeta potentials with single-droplet resolution (Figure 1B-D). μFFE has been previously used for the measurement of protein charge (*31*, *32*) and the separation of proteins and nucleic acids (*33*), and relies on the flow of an analyte through a measurement chamber while an electric field is applied perpendicular to the flow direction. Here, we adapted this technique for single-droplet zeta potential measurements, which allows us to study condensates and their zeta potentials in solution without any surface interactions. The experimental approach is illustrated in Figure 1B–E. After condensates are injected into the μFFE microfluidic device (Figure 1B), they move in response to the applied voltage (Figure 1C, left), and their positions are recorded as a measure of electrophoretic mobility (Figure 1C, right panel). Once positions of individual droplets are quantified from the fluorescence images, the zeta potential can be directly obtained, as further described in Supplementary Materials. In this manner, zeta potential distributions from measurement of thousands of individual condensates can be obtained within a few minutes (Figure 1D). This approach thus allows for the high-resolution quantification of zeta potentials at the single-particle level, which is especially important for samples that are poly-dispersed both in zeta potential and size as is the case for liquid biomolecular condensates.

With the μFFE approach, zeta potentials were acquired for three different biomolecular condensate systems. We first focused on a dipeptide repeat derived from the hexanucleotide repeat expansion in the chromosome 9 open reading frame 72 (*C9orf72*) gene, implicated in amyotrophic lateral sclerosis (ALS) (*34*, *35*). The peptide used consisted of 25 repeats of the dipeptide proline-arginine (PR_25_). This type of peptide is well known to phase separate when mixed with negatively charged polymers (*4*, *35*), including single-stranded RNA consisting of 2500–3500 bases (molecular weight from 800–1000 kDa) of uridine (PolyU). In addition to PR_25_, the protein fused in sarcoma (FUS) was studied. FUS is a widely expressed RNA-binding protein that has been shown to phase separate and has been correlated with ALS phenotypes (*36–38*). We also studied the disease related mutant FUS G156E, which is known to have a faster transition from the liquid-condensed state to the solid state (*10*). Both FUS variants were expressed with a C-terminal EGFP fluorescent protein tag for visualization purposes. The proteins and the peptide:RNA system typify two distinct classes of condensates: those formed via homotypic interactions (*e.g.*, multivalent interactions between the disordered regions and domains of FUS (*39*)), and those sustained by heterotypic interactions (*e.g.*, the association of polyanions and polycations in the PR_25_:PolyU system through complex coacervation (*40*)).

Each of the phase separating systems was assessed using μFFE to determine zeta potential distributions from thousands of individually probed biomolecular condensates. Figure 2 shows the range of zeta potentials obtained across the different protein condensates, as given by their mean values (*μ*), and their degree of heterogeneity, as assessed by the standard deviation of the distributions (*σ*). The trend of absolute zeta potentials of the condensates from largest to smallest was PR_25_:PolyU > FUS wild type > FUS G156E, with mean zeta potential values ranging from –40.6 mV to –15.0 mV. The distributions also showed that the condensates are poly-dispersed in zeta potential, as evident by standard deviations around 11–13 mV. Further analysis revealed that the condensate systems are poly-dispersed in size; yet there is no distinct correlation between zeta potential and size (Figure S3).

**Figure 2.**
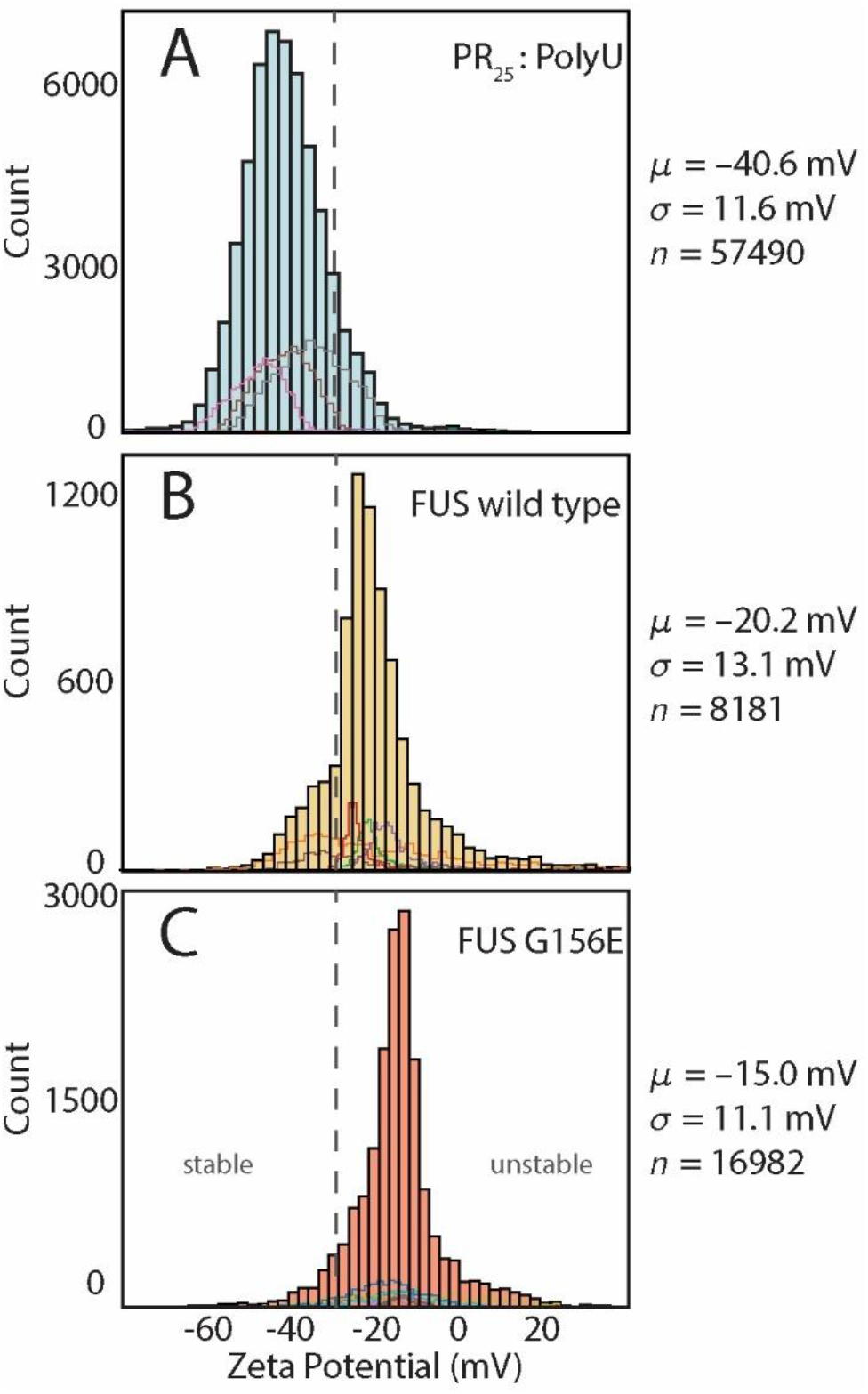
Single-droplet zeta potential measurements of biomolecular condensates. Histograms of single condensate zeta potential measurements for **(A)** PR_25_:PolyU, **(B)** FUS wild type, and **(C)** FUS G156E condensates. Histograms were obtained from all measurements taken on a particular condensate system across all voltages applied, as illustrated in Figure 1. Solid line distributions in each panel at the bottom of each filled histogram represent a collection of measurements from a single replicate at a particular voltage value. Mean, *μ*, and width, *σ*, of distributions as well as number of droplets, *n*, probed are given. Dashed lines indicate boundaries for stable and unstable dispersion with zeta potential cut-offs at −30 mV (*24*, *28*).

### Correlating zeta potential with fusion propensity

The zeta potential is a fundamental parameter to infer the long-range repulsion between colloidal particles in solution and to delineate the stability of emulsions against coalescence or fusion and clustering. Therefore, we hypothesized that the trend in zeta potential values observed for the different condensate systems could reflect their emulsion stability (*i.e.*, their resistance to fuse and coalesce). To test this hypothesis, we assessed the fusion propensity of PR_25_:PolyU and FUS condensates by monitoring droplets merging using light microscopy (*9*, *35*). We observed that PR_25_:PolyU condensates remain stable against fusion (Figure 3A,B), as has been previously reported (*35*). Specifically, PR_25_:PolyU condensates were able to come into contact without fusing, and remain stable over many hours without fusing or clustering. Conversely, FUS wild type condensates rapidly fuse and cluster within seconds to minutes after mixing (Figure 3C,D), in line with previous observations (*10*), and readily exhibit clustering behavior in solution. Similarly, FUS G156E condensates rapidly fuse together within minutes after phase separation (Figure 3E). These observations indeed suggest that there is a correlation between zeta potential and a barrier to condensate fusion.

To corroborate these observations, we further conducted controlled fusion experiments using dual-trap optical tweezers (*10*, *14*) (Figure 4). In these experiments, PR_25_:PolyU condensates showed a higher resistance against fusion compared to FUS wild type condensates. Whereas FUS condensates fused immediately upon contact, PR_25_:PolyU condensates required an additional force to initiate a fusion event, indicating the presence of a repulsion between the condensates. This characteristic is evident in images of moderately deformed PR_25_:PolyU droplets just before fusion, and in the force measurements from optical tweezer experiments (Figure 4A). Here, we observed a dip in the laser signal just before PR_25_:PolyU droplet fusions, indicative of an increased repulsive force between the condensates. This feature was absent in FUS wild type condensates. These observations correlate well with the findings that PR_25_:PolyU condensates have a greater absolute zeta potential compared to FUS, and thus show that a greater absolute zeta potential indeed correlates with an increased barrier to fusion. Interestingly, although there seems to be a higher energy barrier to initiate droplet fusion in PR_25_:PolyU condensates (Figure 4B, bottom panel), once started, fusion proceeds much faster for PR_25_:PolyU condensates than for FUS wild-type condensates (Figure 4B, top panel), suggesting that there is no correlation between the barrier to fusion and the fusion rate.

**Figure 3.**
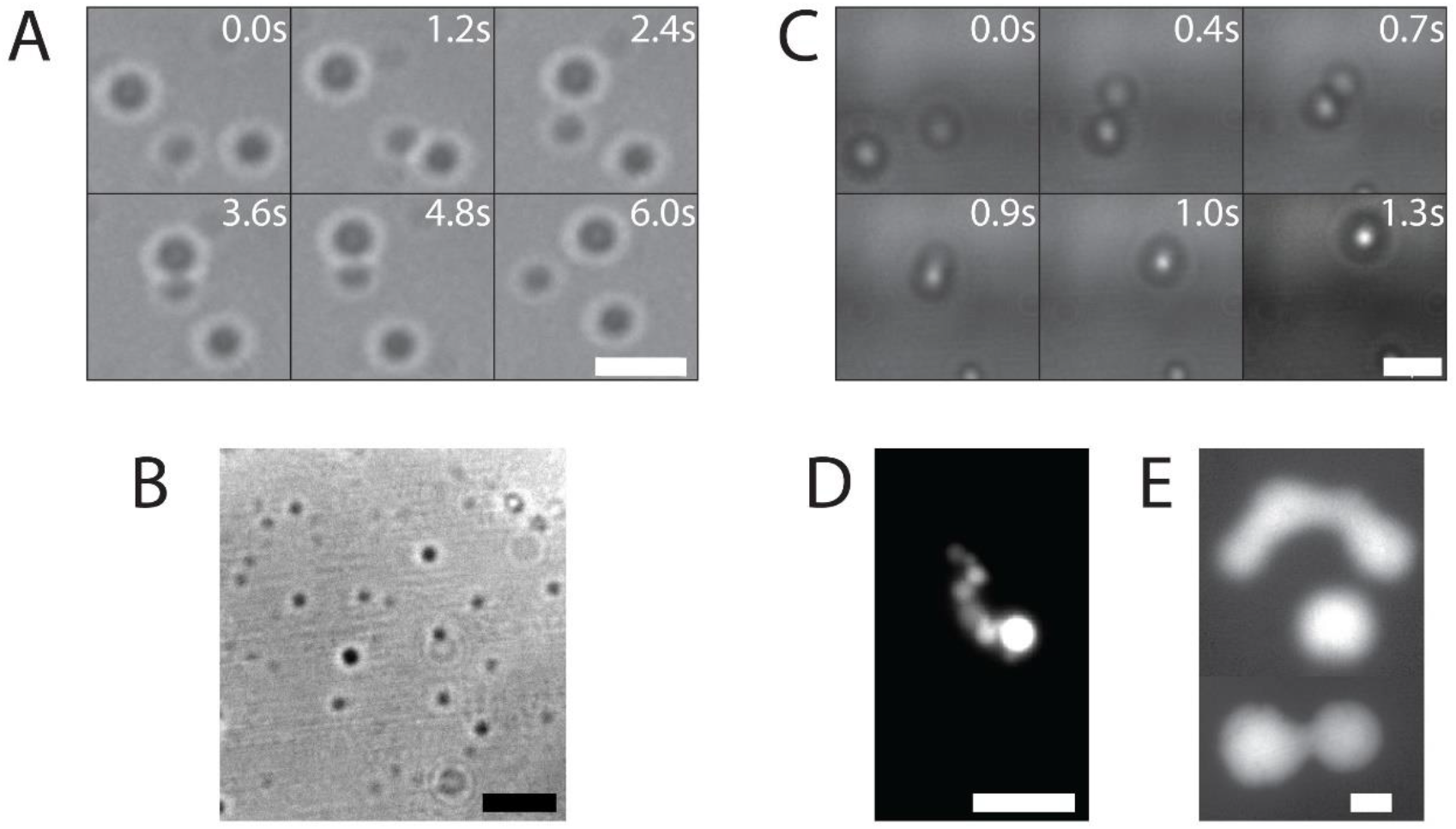
Observations of condensate stability from epifluorescence (Epi) and phase-contrast (PhC) microscopy. **(A, B)** Images of PR_25_:PolyU (PhC), **(C,D)** FUS wild-type (PhC, Epi), and **(E)** FUS G156E (Epi) condensates. All scale bars are 3 μm.

**Figure 4.**
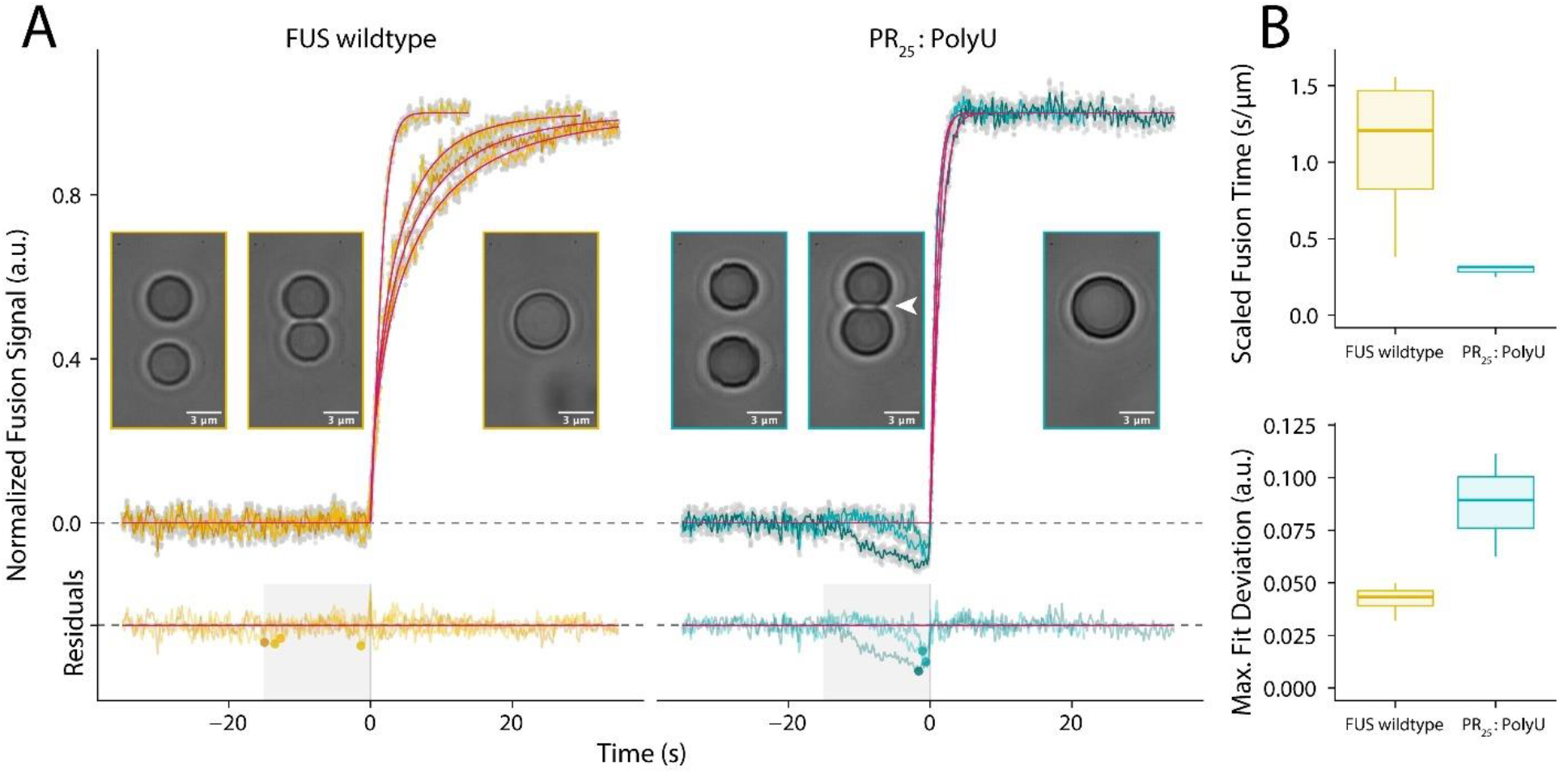
Assessment of condensate stability in controlled coalescence experiments. (**A**) Example traces of controlled droplet fusions using optical tweezers of FUS wild-type and PR_25_:PolyU droplets, together with model fits (magenta) and corresponding residuals. 5% of the raw data (grey points) and smoothed signals (colored lines) are displayed for individual fusion events. For each condition, representative images before fusion, at the onset of fusion, and after fusion are shown. PR_25_:PolyU droplets exhibited a clear indentation (white arrow) before fusion initiated. A significant deviation from the standard fusion model, as illustrated by the dip in the residuals, reflects an energy barrier to be overcome to induce PR_25_:PolyU droplet fusion. We used a window of 15 seconds before fusion onset to quantify the maximum deviation from the model (colored data points). **(B)** Top panel: Size normalized relaxation times indicate that once initiated, PR_25_:PolyU droplets fuse faster than FUS wild-type droplets. Bottom panel: Maximum deviation from the standard model serves as a proxy for the repulsive force required to start fusion.

### Multiscale molecular simulations

To understand the molecular origin of the measured zeta potential values and explore whether or not they correlate with variations in the molecular organization within condensates, in particular the spatial distribution of charged amino acids and the concentration of ions within, we developed a multiscale molecular simulation approach that exploits the advantages of coarse-grained and all-atom models (Figure 5). We started by using a reparameterization of the sequence-dependent LLPS coarse-grained model of the Mittal group (*15*, *41*, *42*) to simulate the formation of FUS and PR_25_:PolyU condensates by means of direct coexistence simulations (*43–45*) of tens to hundreds of interacting biomolecules (Figure 5; Step 1). The reparameterization was implemented to recapitulate the higher LLPS propensity observed experimentally for the full FUS protein versus that of its disordered prion-like domain (PLD) (*14*, *15*, *46*). Subsequently, we performed a back-mapping procedure to convert equilibrium coarse-grained condensates into fully atomistic systems, including explicit solvent and ions (Figure 5; Steps 2–4), and investigated differences in the absorption and distribution of ions between the condensed and dilute phase in both systems (Figure 6). Such a multiscale procedure (Figure 5) is necessary because, on the one hand, investigating the self-organization of proteins into condensed liquids is only feasible with coarse-grained models, given the large system sizes and long timescales required, and on the other hand, capturing changes in counterion behavior requires an explicit all-atom description of biomolecules, water, and ions.

**Figure 5.**
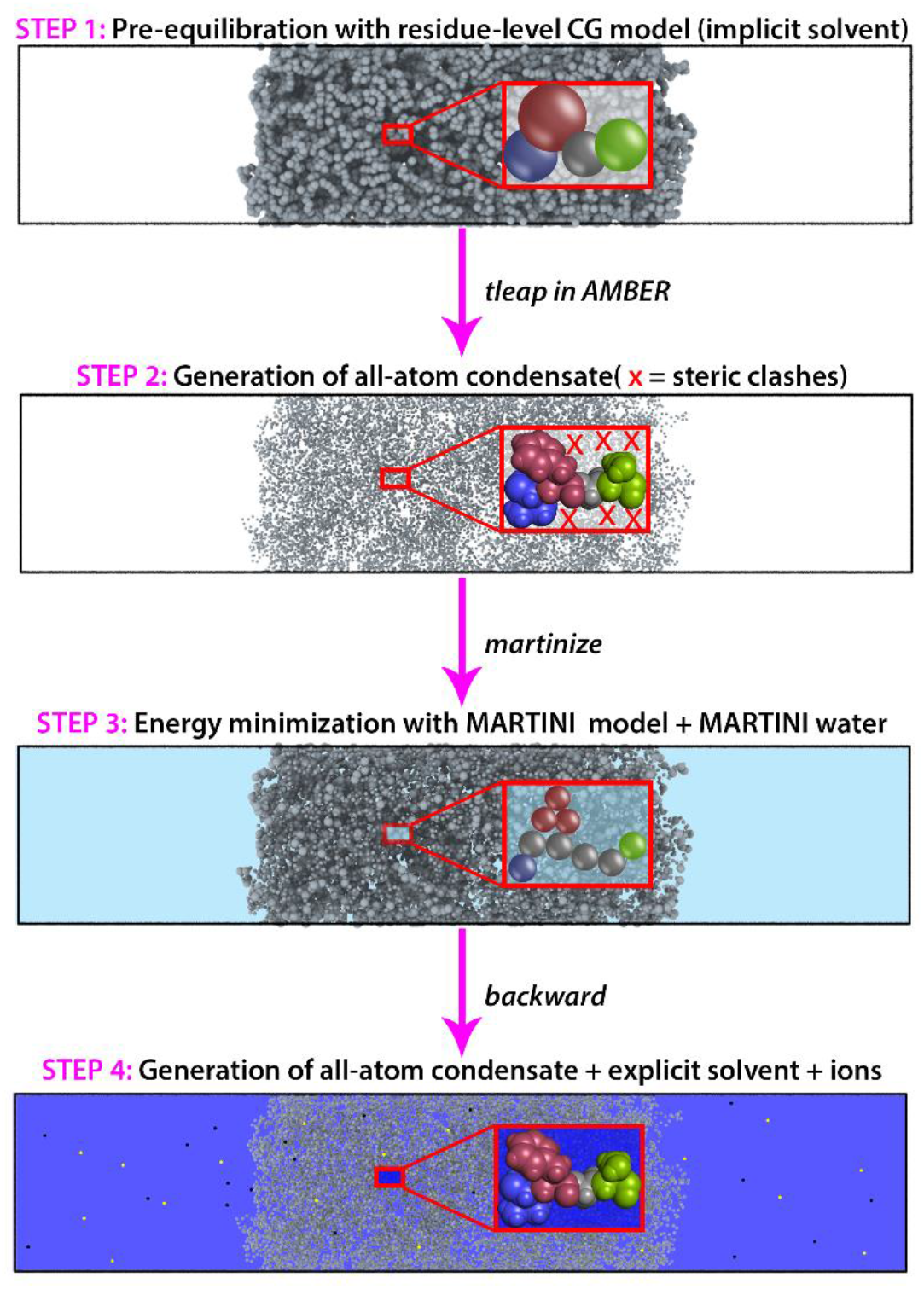
Multiscale strategy for generation of all-atom condensates from pre-equilibrated coarse-grained simulations. ***Step 1***: The system is first equilibrated at residue-level resolution using a reparameterization of the Mittal group coarse-grained model (*41*, *42*). ***Step 2***: The coarse-grained bead coordinates are unwrapped across the periodic boundaries and unwrapped bead positions are defined as coordinates for the amino-acid Cα atoms. Using the tleap module of Amber16 (*71*), missing sidechain and backbone atoms are added in random orientations. ***Step 3***: Because adding atoms in this way results in significant atomic overlaps that cannot be resolved via standard energy minimization procedures, atomistic configurations are mapped to the higher-resolution coarse grained model Martini (*72*) and standard Martini Water (*73*). The system’s energy was then minimized. ***Step 4***: Finally, the program “backward” (*74*) was used to back-map the Martini configuration to the atomistic resolution.

**Figure 6.**
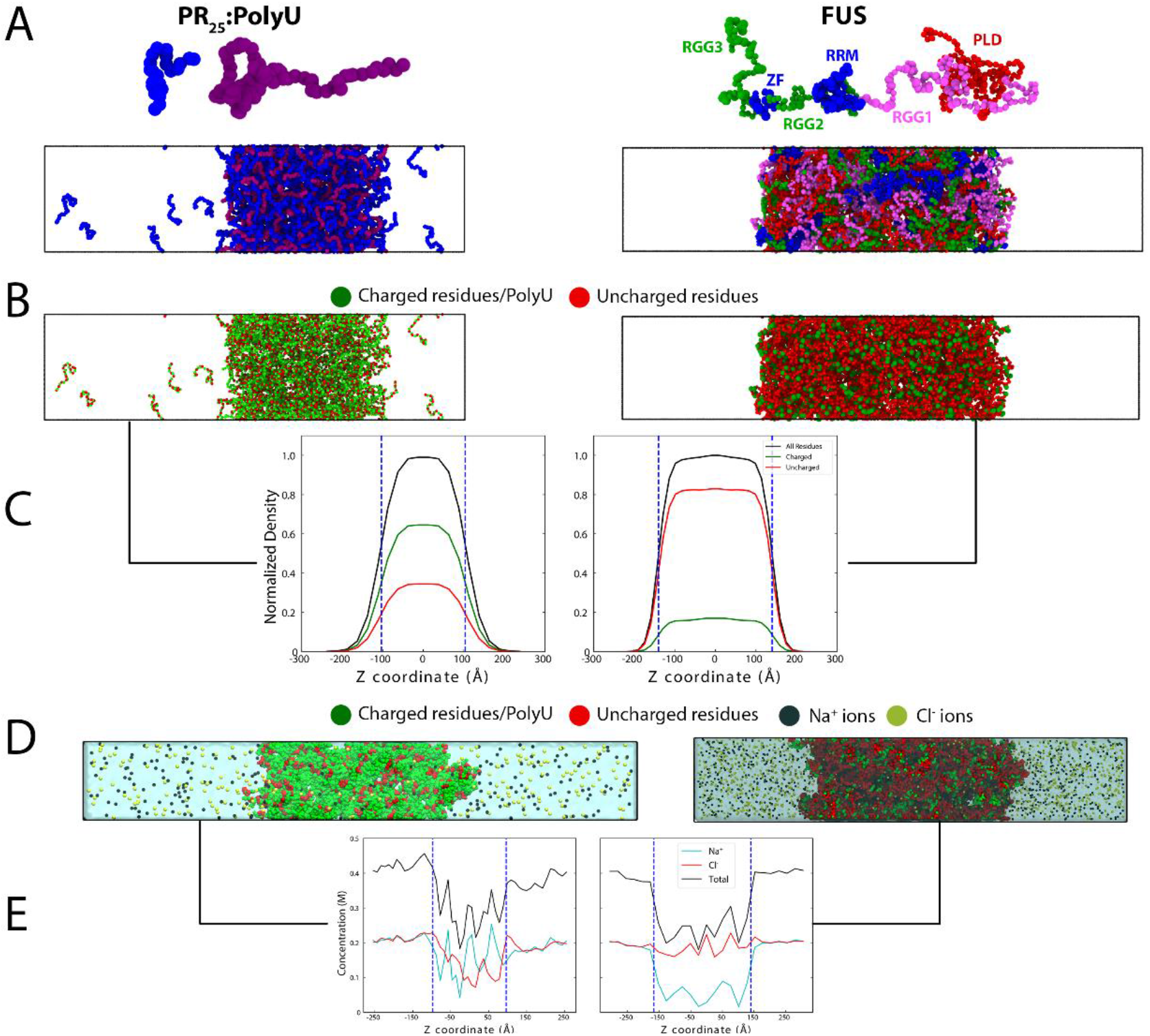
Molecular organization of PR_25_:PolyU and FUS condensates. **(A)** Top panel: One-bead per amino acid/nucleotide coarse-grained representation of PR_25_ (blue), PolyU (purple) and FUS with the PLD (residues 1–165) in red, the extended arginine rich region 1 (RGG1; residues 166–284) in pink, the RNA-recognition motif (RRM; residues 285–371) and the Zinc Finger region (ZF; residues 423–453) in blue, the arginine rich regions 2 (RGG2; residues 372–422) and 3 (RGG3; residues 454–526) in green. Bottom panel: Representative coarse-grained equilibrium configurations obtained via direct coexistence molecular dynamics simulations (*i.e.*, both liquid phases simulated in the same simulation box) of (left) PR_25_:PolyU and (right) FUS condensates. **(B)** Representative configurations from A but with charged species (amino acids and PolyU) colored green and uncharged residues colored red. **(C)** Normalized density of charged and uncharged species across the long side of the simulation box estimated over the coarse-grained equilibrium ensemble showing a much higher concentration of charge in PR_25_:PolyU. The vertical dashed lines show the location of the edge of the condensate. **(D)** Back-mapped atomistic system from equilibrium coarse-grained configuration used to estimate the differential behavior of ions in PR_25_:PolyU and FUS condensates. **(E)** Ion distributions in PR_25_:PolyU and FUS condensates estimated from atomistic direct coexistence molecular dynamics simulations. The vertical dashed lines indicate the approximate locations of the condensate interfaces (condensates are positioned in the center and are in contact with a surrounding diluted phase). The simulations were prepared ensuring similar equilibrium concentrations of ions in the diluted phases of both systems.

Back-mapping coarse-grained protein condensates into all-atom configurations is not a trivial task as it requires using a single bead position for each amino acid to reconstruct them atomistically; that is, by adding all the missing backbone and side chain atoms, while preserving their correct molecular geometry and connectivity, and simultaneously avoiding steric clashes, which are more probable within the crowded environment of the condensates. To tackle this challenge, our multiscale approach is anchored in an innovative back-mapping procedure that breaks the problem down into three simpler steps (Figure 5; Steps 2–4). Each step utilizes standard and widely available biomolecular modelling tools, making our overall procedure easily implementable, widely available, and fully transferable to other condensed-phase protein systems. Accordingly, after equilibrating the protein condensates at coarse-grained residue-resolution (Figure 5; Step 1), we begin the back-mapping by using the coarse-grained bead positions as coordinates for the amino-acid Cα atoms, and add the missing sidechain and backbone atoms in random, and hence potentially spurious, orientations, but maintaining the correct molecular connectivity (Figure 5; Step 2). We achieve this by using the tleap module of Amber16 (19). Adding atoms in this way results in significant atomic overlaps, especially within and close to the more crowded globular regions of multidomain proteins, which cannot be easily resolved through standard energy minimization procedures. To dispose of the numerous steric clashes, in the next step, we coarse-grain the spurious atomistic condensates into the high-resolution Martini model for proteins (20), the ‘soft’ Martini parameters without elastic bonds for PolyU (22), and add standard Martini Water (Figure 5; Step 3). Reducing the resolution back from atomistic to an intermediate coarse-grain level (*i.e.*, between the Mittal group coarse-grained model and the all-atom resolution), we decrease dramatically the number of atoms that incur in steric clashes, while still preserving an explicit representation of backbone and side-chain atoms. This second step is key to make our approach applicable not only to intrinsically disordered peptides but also to large multidomain proteins with globular regions, like FUS. After successfully minimizing the energy at the Martini resolution, in the last step we use the program “backward” (23) to back-map the Martini configuration (discarding the water), now free of atomic overlaps, into full atomistic resolution, and then add explicit water and ions using standard procedures (see Supplementary Materials) (Figure 5; Step 4).

### Insights into the molecular organization of condensates

Our multiscale simulations reveal that both PR_25_:PolyU and FUS condensates (Figure 6A,B) exhibit a mostly homogeneous distribution of charged and uncharged species at physiological salt (Figure 6C). This is not surprising for the highly symmetric and charge-patterned PR_25_:PolyU system, as LLPS here is mainly enabled by electrostatic Arg:U interactions at physiological salt. Indeed, we find a uniform distribution of all species (U, Pro, Arg) at the core of the PR_25_:PolyU condensates (Figure 6; PR_25_:PolyU). More surprisingly, a mostly homogenous molecular organization for FUS condensates is quite remarkable given the molecular complexity of the FUS sequence (see Supplementary Materials). The 526-residue FUS polypeptide chain can be partitioned into an uncharged disordered PLD enriched in Gln, Gly, Ser, and Tyr (residues 1–165), three positively charged disordered Arg-Gly-Gly (RGG) rich regions (RGG1: residues 166–267, RGG2: residues 371–421, and RGG3: residues 454–526), and two globular regions (a RNA-recognition motif: residues 282–371, and a zinc finger: residues 422–453) (*46*). In agreement with experiments (*14*, *47*, *48*), we find that FUS condensates are most strongly stabilized by both electrostatic (*i.e.*, charge–charge and cation–π interactions between the RGG1/3 regions and the Tyr-rich PLD) and hydrophobic (*i.e.*, PLD–PLD) interactions, and more modestly by interactions involving the other domains (Figure S5). Regardless, these preferential patterns of interactions among FUS regions/domains result in homogeneous condensates in the conditions we probed.

A crucial difference between the molecular organizations of FUS and PR_25_:PolyU condensates is the much higher concentration of charged species (both positive and negative) in PR_25_:PolyU condensates versus FUS, including those at the condensate surfaces (Figure 6C and 6E, and Figure S6). Importantly, although the core of PR_25_:PolyU condensates have a homogeneous distribution of positive and negative molecules, the surface itself is more concentrated in PR_25_ peptides (Figure S7), and hence is rich in positive charge. We hypothesize that preferential positioning of PR_25_ peptides towards the interface stems from the lower valency of such molecules in comparison to that of the much longer polyU polymers; this is because concentrating lower valency species, that sustain fewer LLPS-stabilizing interactions, towards the interface is expected to minimize the interfacial free energy of the condensate (*49*). Importantly, when me measure the interfacial free energy of the condensates in our simulations, we find that its value for FUS condensates (0.35±4 mJ/m^2^) is almost twice of that for PR_25_:PolyU droplets (0.20± 4 mJ/m^2^). We note that the difference we observe is qualitative as the coarse-grained simulation do not include explicit solvent, and thus likely underestimate the absolute value of the interfacial free energy in both condensates. Despite this, the trend is that condensates which concentrate more charged species (e.g., positively charged PR_25_ tails) at the surface (Figure S7) tend to have lower interfacial free energies. Concomitantly, concentrating positively-charged PR_25_ tails at the surface results in a higher electrostatic repulsion among individual PR_25_:PolyU condensates, than that among FUS condensates, which shows a more electroneutral surface. Both effects, lower interfacial free energy and higher electrostatic repulsion, challenge droplet fusion in PR_25_:PolyU condensates. These observations are in full agreement with the larger absolute zeta potential values we measured experimentally for PR_25_:PolyU compared to FUS.

Besides a notably higher density of charged species (Figure 6B,C; left), PR_25_:PolyU condensates establish more favorable electrostatic interactions with counterions than FUS condensates. The high concentration of charge at PR_25_:PolyU surfaces is evident from the higher density of counterions at the interface than at the condensate core, and most notable of Cl^−^ ions, which are needed to screen the solvent-exposed PR_25_ tails. In agreement, FUS condensates, which contain less charged amino acids overall (Figure 6B,C; right), also absorb a lower total concentration of counterions. Indeed, because FUS is almost fully devoid of negatively charged residues, Na^+^ is present at very low concentrations inside FUS condensates.

As expected, our simulations reveal that counterions slow down (*i.e.*, have a smaller diffusion coefficient) when they enter the condensed phase, where they find many kindred species to bind transiently to (*50*). Interestingly, counterions diffuse more slowly within FUS condensates than within PR_25_:PolyU condensates (Table S1). This observation likely stems from the higher molecular density of FUS condensates (~0.54 g/cm^3^) versus PR_25_:PolyU condensates (~0.40 g/cm^3^), the abundance of Arg residues in FUS available to establish strong cation-anion interactions with Cl^−^, and the lack of other negatively charged species to displace Cl^−^ from their FUS absorption sites. Consistently, in the more charge-rich PR_25_:PolyU condensates, counterions diffuse slightly more freely because of the lower condensate density, and since Arg and U are already paired up and establish strong cation– anion interactions (Table S1).

Collectively, our simulation results and experimental zeta potential measurements suggest that larger absolute zeta potential values occur in systems that are more highly charged overall, and, importantly, that exhibit a higher total charge at the surface. In such systems, LLPS is usually more heavily driven by electrostatics suggesting that larger absolute zeta potential values correlate with stronger and longer-range intermolecular interactions within condensates.

The wide variations in condensate zeta potential values measured experimentally are indicative of surface heterogeneity, both in shape and charge distribution. This notion is supported by our simulations, which reveal a highly dynamical behavior of biomolecules inside condensates, especially at the interfaces, where a continuous dynamical reconstruction of the interfacial structure occurs via capillary wave fluctuations (*51*, *52*). Biomolecules within liquid condensates sample a wide range of conformations and interconnect with one another through weak short-lived bonds; this phenomenon is intensified at the droplet boundaries, where proteins are less favorably solvated. The continuous dynamical rearrangement of the interface, including protein exchanges from both phases, which induces charge and geometric droplet heterogeneities, is consistent with the heterogeneous zeta potential values we measure. Additionally, it is likely that the smallest droplets within the polydisperse condensate distribution are affected by curvature effects (*53*, *54*), inducing variations in the droplet surface tension as a function of their size. Since the stiffness of the interface is directly related to the droplet surface tension, such variations might result in even more heterogenous capillary wave profiles (*55*).

## Discussion

Through the development of a μFFE approach for probing electrophoretic properties of phase-separated condensates, we were able to quantify the zeta potential of biomolecular condensates with single-droplet resolution and correlate this parameter to condensate stability against fusion. Our results show that PR_25_:PolyU condensates have a higher absolute zeta potential than FUS wild type and G156E mutant condensates, and this trend correlates well with qualitative emulsion stability observations from microscopy experiments and quantitative data from optical tweezer measurements. Through multiscale molecular dynamics simulations, we show that the differences in absolute zeta potential values, and hence the stability of biomolecular condensates against fusion, emerges from distinctly different molecular organizations of the condensates. While PR_25_:PolyU condensates are stabilized mostly by electrostatic interactions and possess highly positively-charged surfaces, FUS droplets are predominantly sustained by cation–π and hydrophobic interactions, and exhibit only modestly charged interfaces. These findings, therefore, establish the surface charge density of condensates as the molecular origin of the modulation of their propensity to fuse, and the zeta potential as a fundamental quantity to infer it. Specifically, we reveal that condensates with more densely charged surfaces, and hence higher absolute zeta potentials, exhibit a higher stability against fusion (*i.e.*, a higher force is needed to induce fusion due to their lower surface tension) and therefore higher inter-condensate electrostatic repulsion. This is consistent with previous studies suggesting that the surface charge of emulsions can have a direct effect on the surface tension (*56*, *57*).

When exposed to the non-equilibrium environment of a cell, droplet stability against fusion may be regulated by additional factors, such as chemical reactions that dynamically alter the concentrations of biomolecules and the chemical compositions in and out of condensates, temperature gradients that impact the relative strength of protein-protein interactions, and concentration gradients (*21*, *58*). Regardless of this, our work provides fundamental molecular information to understand one of the mechanisms by which such additional non-equilibrium process might modulate droplet emulsion stability, namely, active control of condensate surface charge. Further, both FUS (*10*, *48*) and PR_25_ (*35*, *59*) have been shown to stabilize condensates *in vitro*, in conditions of thermodynamic equilibrium (*i.e.*, in the absence of additional active or catalytic processes). Our work proposes that such passive stabilization stems from a combination of repulsive forces between condensates, and the effects that surface electrostatics have on lowering the surface tension of the droplets.

The correlation between electrostatic properties and condensate stability against fusion, as predicted by classical emulsion theory, provides a means by which protein condensates can be classified and compared according to their zeta potential (*25*, *26*). A larger absolute value of zeta potential confers greater resistance against coalescence and clustering, as has been shown for various oil-in-water emulsions of phosphorylated species and poly-amino acid stabilized inorganic emulsions (*27–29*). Moreover, a threshold of 30 mV in absolute zeta potential seems to exist for biomolecular condensates, as has been previously put forward in literature (*24*, *28*), above which higher stability against coalescence is observed and below which condensates show increased propensity for clustering and coalescence. Along with this cut-off, the significant variability in zeta potential, evident by the wide distributions, indicates that a single ensemble of condensates will have varied degrees of fusion propensities within it. The measurement of zeta potentials also revealed that the surfaces of condensates possess markedly different surface charges. Our multiscale molecular simulations further demonstrate that proteins positioned at the condensate interface, which we anticipate impact most significantly the zeta potential values, have a higher tendency to dynamically transition in and out of condensates. That in turn is expected to alter the structure and properties of the droplet interface and, hence, the exact value of the total charge surface of the condensate, explaining the heterogeneity in zeta potential values. These results offer a deeper understanding of the internal and surface geometry of condensates.

A further observation is that the zeta potentials of biomolecular condensates can considerably vary even though their overall composition remains constant. This variation is indicated by the large standard deviations for the zeta potential distributions, which ranged from 24% to 78% of the mean, while the standard errors were all well below 0.1% due to the large sample size. Thereby, further suggesting that heterogeneity with respect to surface geometry is present across condensates within a single sample, which may be partly due to dynamic rearrangement and exchange of proteins both within the condensates and with the exterior. This constant reassembly is consistent with the description of condensates as highly dynamic assemblies (*37*, *60*). Our multiscale simulations also reveal the highly dynamical nature of the condensates. Specifically, biomolecules adopted diverse conformations and, due to weak intermolecular interactions, dynamically switched their interaction to other neighbors. In some cases, proteins even escaped to the diluted phase and were subsequently recruited back into the condensate, thus changing the shape and chemical composition of the interface continuously. Indeed, the ability of biomolecules to form many weak interconnections within a random and dynamical percolated network is critical to the stability of biomolecular condensates (*49*).

The observed correlation between emulsion stability and zeta potential also has important implications for diseases, specifically for the transition of condensates from their liquid state to solid aggregates. It has been shown that FUS can transition into toxic aggregates associated with the onset and development of motor neuron disease more readily when it is contained in condensates (*10*), and this trend holds true for other proteins as well, including TDP-43 and other condensate forming systems (*61*, *62*). Recent theoretical work has highlighted how condensates could behave as compartments for aggregate formation, and has also indicated how more aggregates could form within condensates of greater size (*63*). In addition, it is well known that the primary nucleation of solid phases is directly dependent on the number of available precursor monomer protein molecules (*64*) and, since monomer concentration is higher in condensates than in the dilute phase (*65*), condensates may serve as epicenters for the formation of toxic solid aggregates. Hence, the propensity of FUS condensates to fuse more readily, as dictated by a low absolute zeta potential, causes the condensates to grow bigger over time in a fusion growth model. In an Ostwald ripening model where growth occurs through transfer of monomer from one condensate to another, a low surface charge also allows for greater growth due to the lack of electrostatic repulsion against incoming monomer (*66*). A larger size may thus render condensates more favorable for nucleation and growth of aggregates. Furthermore, the lower absolute zeta potential observed for FUS G156E compared to FUS wild type might explain its higher propensity to form aggregates (*10*).

Beyond pathophysiological implications, the immiscibility and size control of phase separated condensates has been indicated to be relevant particularly in the control of the size of organelles during cell growth and embryonic development (*67*, *68*). Additionally, the size of condensates could be a marker of cancer proliferation (*69*), suggesting that the modulation of the size of certain condensates, controlled through their zeta potential, could be exploited for therapeutic interventions. In contrast, the zeta potential of condensates can also provide information regarding their propensity to fuse under physiological conditions, such as is the case for nucleoli which are able to organize RNA due to their fusion in a single large condensate(*20*).

Taken together, this work establishes the zeta potential as a fundamental quantity to infer the emulsion stability of biomolecular condensates, and proposes a transferable multiscale molecular approach to connect mesoscale properties of condensates to the atomistic properties of the proteins that are contained within them. By probing the zeta potential on a single condensate level, we described the electrostatic nature of PR_25_:PolyU and FUS condensates and correlated these experimental results with their observed stability from fusion experiments. Our multiscale molecular approach further described the detailed molecular behavior of these condensates, including their surface charge density and its impact on their interfacial free energy, the intermolecular interactions of the component biomolecules, and the distribution and mobility of ions in- and outside of condensates. Overall, these results expand our understanding of the physical and molecular factors that control the emulsion stability of condensates.

## Materials and Methods

### Materials

All reagents and chemicals were purchased with the highest purity available. The PR_25_ peptide, containing 25 proline–arginine repeats, was obtained from GenScript. N-terminally labelled PR_25_ was obtained by reacting the peptide with amine-reactive AlexaFluor546 (Sigma-Aldrich). PolyU RNA with a molecular weight range from 800– 1,000 kDa was purchased from Sigma-Aldrich. FUS wild type and FUS G156E were produced as C-terminal EGFP fusion proteins as previously described (*10*) and stored in 50 mM Tris-HCl (pH 7.4), 500 mM KCl, 1 mM dithiothreitol, 5% glycerol. PR_25_ phase separation was induced by mixing 100 μM PR_25_ peptide with 1 mg/mL PolyU RNA in 5 mM Tris-HCl (pH 7.4). For both FUS variants, phase separation was induced by diluting the proteins to a final protein concentration of 3 μM in 25 mM KCl, 5 mM TRIS (pH 7.4). For PR_25_:PolyU and both FUS mutants, the phase separated condensates were analyzed via μFFE within ~10 min of creation in order to minimize ageing effects; no systematic differences in zeta potential were observed across replicate samples on this time scale. 60 nm fluorescently labelled spherical gold nanoparticles (NanoPartz) were used for control measurements mentioned in Figure S4.

### μFFE experiments

The design of the 3D μFFE microfluidic chip with liquid electrodes was adapted from a device previously used for studying protein charge and the separation of biomolecules (*31*, *33*). A schematic is shown in Figure S1. The device, constructed from a top and a bottom layer, was fabricated using standard single- and multilayer photolithography techniques as described in detail in the Supplementary Materials. Briefly, the microfluidic channels within each layer were patterned into polydimethylsiloxane (PDMS; Sylgard184, Dow Corning) using SU-8 photoresist (Microchem) on silicon masters (MicroChemicals). Top and bottom PDMS layers were then connected through plasma bonding and subsequently bonded to glass microscope slides using oxygen plasma (Diener Electronics). Devices were operated as detailed in the Supplementary Materials and fluids introduced using automated syringe pumps (neMESYS, Cetoni). Electric potentials were applied using a programmable 500 V power supply (Elektro-Automatik EA-PS 9500-06) and images acquired using a Zeiss AxioObserver D1 microscope. Further details are given in the Supplementary Materials. Image and data analysis were performed using the Fiji/ImageJ data processing software and custom-written Python scripts, respectively. Zeta potentials were calculated as described in detail in the Supplementary Materials.

### Epifluorescence and phase-contrast microscopy in droplet stability experiments

For experiments assessing condensate emulsion stability, epifluorescence and phase contrast images were captured using an AxioObserver D1 microscope (Zeiss) with either a 40x or 100x air objective after the specified aging time for each sample (Figure 3). Condensates were imaged within a 50 μm tall microfluidic imaging chamber in the same buffer conditions as utilized for μFFE experiments.

### Optical tweezer measurements

Condensates were phase-separated in 5 mM Tris-HCl, 25 mM KCl, pH 7.4 and immediately applied to a sample chamber. Two droplets were trapped in two optical traps of the same trap stiffness. With the first trap stationary, the second trap was moved to bring the droplets into contact and initiate fusion. If fusion did not occur upon first contact as in the case of PR_25_:PolyU condensates, the second trap was further moved to push the droplets together. As soon as coalescence initiated, the traps were kept stationary. Laser signals were recorded at 1 kHz resolution. Signals from the two traps, equal in magnitude and opposite in sign, were combined into the differential signal, from which coalescence relaxation times were deduced. A random sample of 5% of the recorded data is plotted as grey points in Figure 4. Raw data were smoothed with a Savitzky-Golay filter of 3^rd^ order and a window of 501 points.

### Fit of optical tweezer traces

The standard model for droplet fusion is based on the assumption that droplets start to coalesce as soon as their surfaces touch. This assumption holds true for many purified protein liquids (*10*, *13*, *14*, *70*). To characterize fusion dynamics, time traces of the tweezer signal, *S*(*t*), were fitted with a stretched exponential model as described previously (*14*). Briefly, the model is defined as:

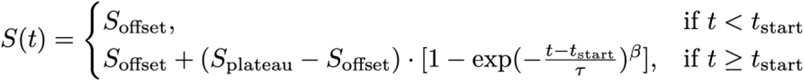

where *τ* denotes the relaxation time, *β* the stretch exponent, *t*_start_ the onset of fusion, *S*_offset_ the signal offset on the detector, and *S*_plateau_ the final signal value after coalescence finished. All fusion traces (Figure 4A) have been normalized and aligned according to the start time of coalescence as deduced from the fit. Residuals from the fit were calculated for the smoothed signal. We took the maximum negative deviation from the standard model within a window of 15 seconds before the onset of fusion as a proxy for the additional energy barrier to be overcome. To quantify the fusion dynamics, the mean relaxation time was normalized by the geometric radius of the two fusing droplets.

### Multiscale molecular simulations

To investigate the molecular organization of proteins, PolyU and ions within the condensates, we develop a two-step multiscale molecular simulation method. The first step consists of coarse-grained molecular dynamics simulations of tens to hundreds of biomolecules to investigate the equilibrium ensembles of FUS and PR_25:_PolyU condensates (see further coarse-grained simulation details in the Supplementary Materials). During the second step, we undertake a back-mapping procedure and perform atomistic molecular dynamics simulations with explicit solvent and ions to assess the distribution of ions in the condensed and diluted phases, and obtain magnitudes directly related to zeta potentials estimations (see details of atomistic simulations in the Supplementary Materials).

## General

We thank members of the Knowles laboratory for discussions. We thank Jeetain Mittal and Gregory L. Dignon for invaluable help with the implementation of their sequence-dependent protein coarse-grained model in LAMMPS.

## Author contributions

T.J.W., G.K., and T.P.J.K. conceived of the idea and designed research. T.J.W. carried out all microfluidic experiments and analyzed the data. J.R.E., J.A.J., A.S., and R.C.G. designed the simulations. J.R.E. and J.A.J. performed the coarse-grained simulations. A.S. performed the atomistic simulations. J.R.E., J.A.J., and A.S. analyzed the simulation results. M.J., T.J.W., and G.K. performed and analyzed optical tweezer experiments. R.C.G. supervised the simulation work. W.E.A. and K.L.S. provided microfluidics chip designs and data analysis approaches. S.A. provided protein materials. T.J.W., G.K., R.C.G. and J.A.J. wrote the manuscript. All authors edited the manuscript.

## Supplementary Materials

### Supplementary Materials and Methods

#### Design of the μFFE device

The design of the μFFE microfluidic chip with liquid electrodes was adapted from a device previously used for studying protein charge and the separation of biomolecules (*1*, *2*). A schematic is shown in Figure S1. The device is 90 μm tall in the central electrophoresis chamber and 5 μm tall in the sample injection port. In total dimensions, the device is approximately 7 mm long and 2 mm wide. The 3D design was utilized to minimize the effect of velocity differences within the channel; further details on device design optimization are given in Saar *et al.* (*1*). For operation, the sample of interest containing phase-separated droplets is flown into the device by the central injection port where it is then surrounded by the carrier buffer solution, which was 5 mM Tris-HCl (pH 7.4) in experiments with PR_25_:PolyU and 5 mM Tris-HCl (pH 7.4), 25 mM KCl in experiments with both FUS variants. On either side of the main channel, liquid electrolyte channels are filled with a constant flow of a 3 M KCl solution, supplemented with 1 mg/mL fluorescein (Sigma-Aldrich) for visualization purposes. The electrolyte solution enters the main channel via 40 μm wide and 5 μm tall electrolyte ridges, which allows for a narrow stream of electrolyte to coat both sides of the main electrophoresis channel. This solution remains under constant flow and acts as a liquid electrode, which is continually replaced. Utilization of liquid electrodes allows for high voltages to be applied as gaseous electrolysis products are flushed out of the device through the hollow electrodes (*3*). Further, the flow of electrolyte also aids in suppression of Joule heating within the device, which can be an issue with other types of micro-scale electrophoresis devices (*4*).

**Figure S1.**
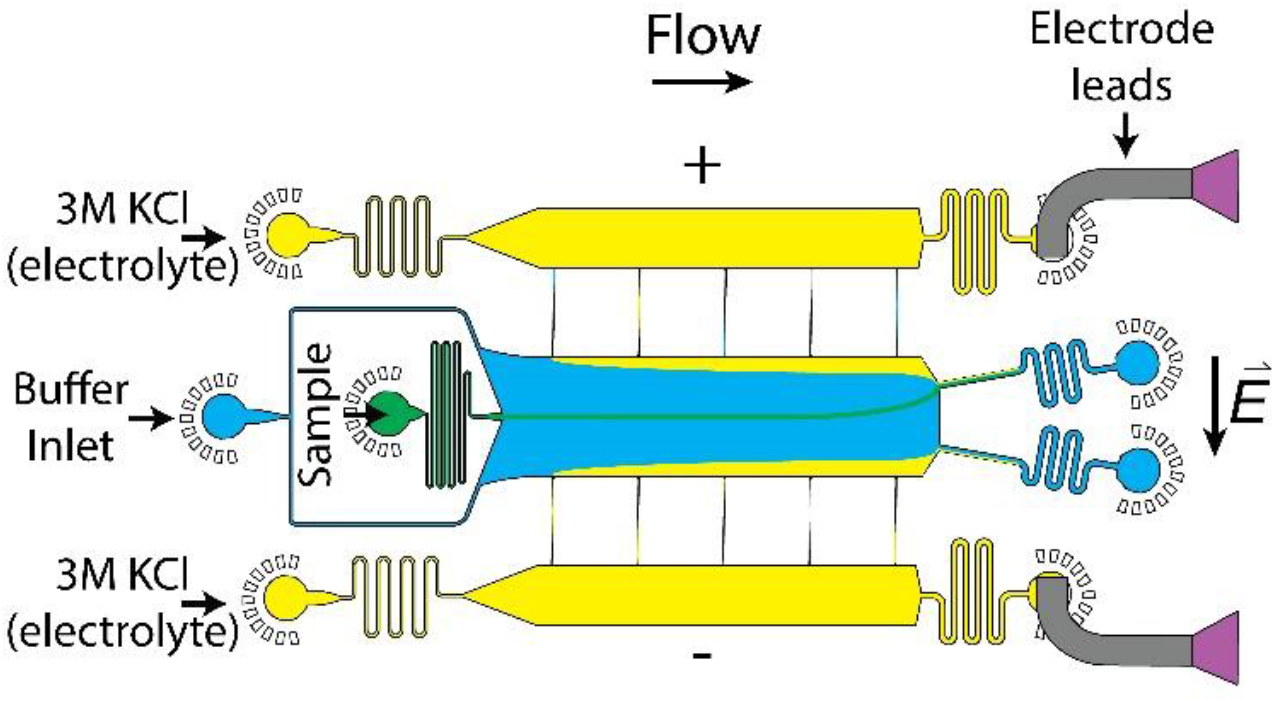
Full schematic of μFFE device. This schematic shows the general design of the 3D μFFE device. The sample is injected through a central port at the beginning of the channel, which is 5 μm tall. Thereby, the sample does not come into contact with any surfaces of the central channel containing co-flow buffer, which is 90 μm tall. The 3 M KCl solution acts as an electrolyte and flows along the edges of the central channel to allow the voltage to be transmitted from the outlet ports to the sample where an electric field is induced opposite to the direction of flow. Further description of the usage and design of the device is given in the text.

#### Fabrication of the μFFE device

Microfluidic masks were first designed using AutoCAD (Autodesk) and desired device geometries then printed on acetate transparencies (Micro Lithography Services). Polydimethylsiloxane (PDMS; DowCorning) devices were produced from SU-8 (MicroChem) molds fabricated via standard photolithographic processes by plasma bonding two individual PDMS chips to each other. Accordingly, two molds were made in order to comprise the two separate sides of the 3D microfluidic devices, with the bottom layer being produced from a single-layer (SL) replica mold, while the top layer was produced from a two-layer (TL) replica mold. Specifically, the mold for the SL chips was fabricated to a height of 45 μm and included all the structures of the devices with the exception of the protein inlet and the electrolyte bridges connecting the electrophoresis chamber and the electrolyte channels. This was achieved by spinning SU-8 3050 photoresist onto a polished silicon wafer (MicroChemicals) followed by standard soft-lithography procedures (*5*) using a custom-built LED based apparatus for performing the UV-exposure step (*6*). The fabrication of the TL replica mold for the top layer involved two subsequent lithography steps performed with SU-8 3005 and 3050 to obtain 5 and 45 μm high channels, respectively. The protein inlet as well as the connecting electrolyte bridges were featured only on the 5 μm layer, while the buffer inlet, the electrophoresis chamber, and the electrolyte channels were fabricated onto the 45 μm layer only are identical to how they appear on the SL replica mold. Feature heights on the master were assessed using a profileometer (DektakXT, Bruker). The top and bottom layer replica molds were then used to fabricate PDMS chips employing a 10:1 prepolymer-PDMS-to-curing-agent ratio (Sylgard 184, DowCorning). After degassing and curing for 3 h at 65°C, the two halves of the devices were then cut out of the molds, and holes for tubing connection (0.75 mm) and electrode insertion (1.5 mm) were created in the top layer PDMS half. Both sides of the devices were cleaned by application of Scotch tape and sonication in isopropanol. Following treatment using an oxygen plasma oven (Femto, Diener electronic) at 40% power for 30 s, the PDMS bottom layer was bonded on a glass slide with the channels facing upward. The PDMS top layer was then placed on top and carefully aligned to create a 3D device. The device was baked at 65°C for 24 h to ensure optimal bonding. Before use, devices were rendered hydrophilic via prolonged exposure to oxygen plasma (500 s, 80% power) (*7*). After this treatment, surface hydrophilicity was prolonged by immediate filling of device channels with deionized water using gel-loading tips (Fisherbrand).

#### Device operation and experimental conditions in μFFE experiments

The device was operated by injecting the sample solution, the carrier buffer solution, and the electrolyte solution into the corresponding inlets using automated syringe pumps (neMESYS, Cetoni). The sample was introduced from a 100 μL glass syringe (Hamilton), other solutions were flowed from 10 mL plastic Norm-Ject syringes (Henke-Sass Wolf). All fluids were introduced to the device by 0.012X0.030” PTFE tubing (Cole-Parmer). Typical values for the flow rates were 5 μL/hr for the sample, 400–500 μL/hr for the carrier medium, and 100– 250 μL/hr for the electrolyte solutions. Fluid waste was guided out of the device by tubing inserted into device outlets. Electric potentials were applied using a programmable 500 V power supply (Elektro-Automatik EA-PS 9500-06) via bent hollow metal dispensing tips (15G, Intertonics) inserted into the electrolyte outlets. The voltage was varied in linear steps, typically in the range between 0 to 80 V, using a computer controller (Raspberry Pi). Simultaneously, current readings using a digital multimeter (34401A, Agilent Technologies) were taken. Schematics of the electrical setup can be seen in Figure S2. The measurements for determining the electrical resistance of the electrodes and estimating the effective electrical potential applied across the devices were performed in an identical manner but with the sample and carrier medium replaced with 3 M KCl solution as has been described in detail earlier. All measurements were performed at room temperature.

#### Optical detection in μFFE experiments

Images were acquired using an inverted fluorescence microscope (Zeiss AxioObserver D1) equipped with a high-sensitivity electron-multiplying charge-coupled device (EMCCD) camera (Evolve 512, Photometrics). In experiments with FUS, an appropriate filter set for EGFP detection was used (49002, Chroma Technology). Exposure times were around ~10 ms for each image, allowing for between 30–100 particles to be imaged per frame, and 500–2000 to be imaged per experimental μFFE run. Due to high amounts of free PR_25_ monomer in solution, images for the PR_25_ system were captured in bright-field mode with a phase contrast ring (Ph2). The movement of the droplets in the microfluidic chip was collected by running samples containing the phase-separated droplets into the main chamber of the device and taking images approximately at the coordinate corresponding to the 4^th^ electrolyte bridge (*i.e.*, approx. after 4 s of travel within the chip). At each voltage, a series of images were taken in order to detect ~500–2000 droplets.

#### Data analysis and calculation of zeta potentials

Images taken in μFFE experiments were analyzed using the Fiji/ImageJ data processing software. Condensates were detected using the TrackMate package (*8*), which returned the x,y-coordinates of individual droplets within the channel, with x being the coordinate in the direction of the length of the channel (*i.e.*, flow direction) and y being the coordinate in the direction of its width (*i.e.*, perpendicular to the flow). By calibrating the position of the image within the channel, the travelled distance in x,y-direction over a stream of images was determined, which subsequently gave the residence time, *t*_r_, needed for drift velocity calculations (*i.e.*, the lateral and longitudinal movement of droplets in time). Accordingly, the drift velocity, *v*, was calculated from the vertical displacement of each condensate, referred to as Δ*y*, according to

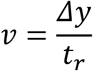

Δ*y* is quantified as the vertical displacement of each condensate from the average vertical coordinate of the stream at 0 V, and *t*_r_ was calculated from the flow rate, the x coordinate (or distance traveled), and the known dimensions of the channel. Note, given that the sample stream height is <5% of the height of the total channel and the co-flow buffer flow rate is 50 times higher than the sample flow rate, not much broadening of the signal from the parabolic flow profile is to be expected, which would occur mainly near the edges of the device and may cause velocity variations across the channel.

With *v* at hand, the electrophoretic mobility, *μ,* was calculated as

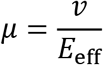

where *E*_eff_ is the effective electric field across the main electrophoresis channel. *E*_eff_ is equivalent to *V*_eff_/*w*, with *V*_eff_ being the effective voltage and *w* being the width of the device, and was obtained through calibration of each device with 3 M KCl as shown in Figure S2B. In order to determine *V*_eff_, first the resistances *R* were determined according to Ohm’s law *R = V/I* for each point shown in Figure S2B. By filling the device with 3 M KCl, the internal resistance is effectively zero; therefore, the resistance of the electrode, *R*_elect_ = *V*_app_/*I*, could be determine from the 3 M KCl calibration measurement. Similarly, the resistance of the entire device could be determined during the sample measurement according to the relation, *R*_dev_ = *V*_app_/*I*.

With the resistances *R*_elect_ and *R*_dev_ at hand, the resistance of only the internal measurement chamber could be calculated as *R*_main_ = *R*_dev_ − *R*_elect_. Thereby, the voltage drop within the main chamber, expressed as a percentage drop could be calculated as the ratio *effV* = *R*_main_/*R*_dev_. Typically, electrical resistances of 115 and 100 kΩ were determined for *R*_dev_ and *R*_elect_, respectively, and we obtained voltage efficiencies varying from 2% to 12%. From this, *V*_eff_ could be calculated according to *V*_eff_ = *effV* × *V*_app_, where *V*_app_ is the applied voltage at the respective sample measurement. This allowed *E*_eff_ to be determined and therefore the mobility *μ* of the droplets to be calculated as described.

The measured electrophoretic mobilities for each condensate could then be converted into the zeta potential, *ζ*, according to the following relation using a modified version of Henry’s function (*9*)

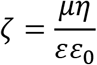

In this relation, *ε* is relative permittivity of the solution, *∊*_0_ is the permittivity of a vacuum, and *η* is the dynamic viscosity of the solution. The solution was treated as water, thus the accepted value of *ε* = 78.5 (*10*) and *η* = 1.0518×10^−3^ Pa s (*11*) were used. All calculations were all carried out in Python using the integrated development environment Spyder.

**Figure S2.**
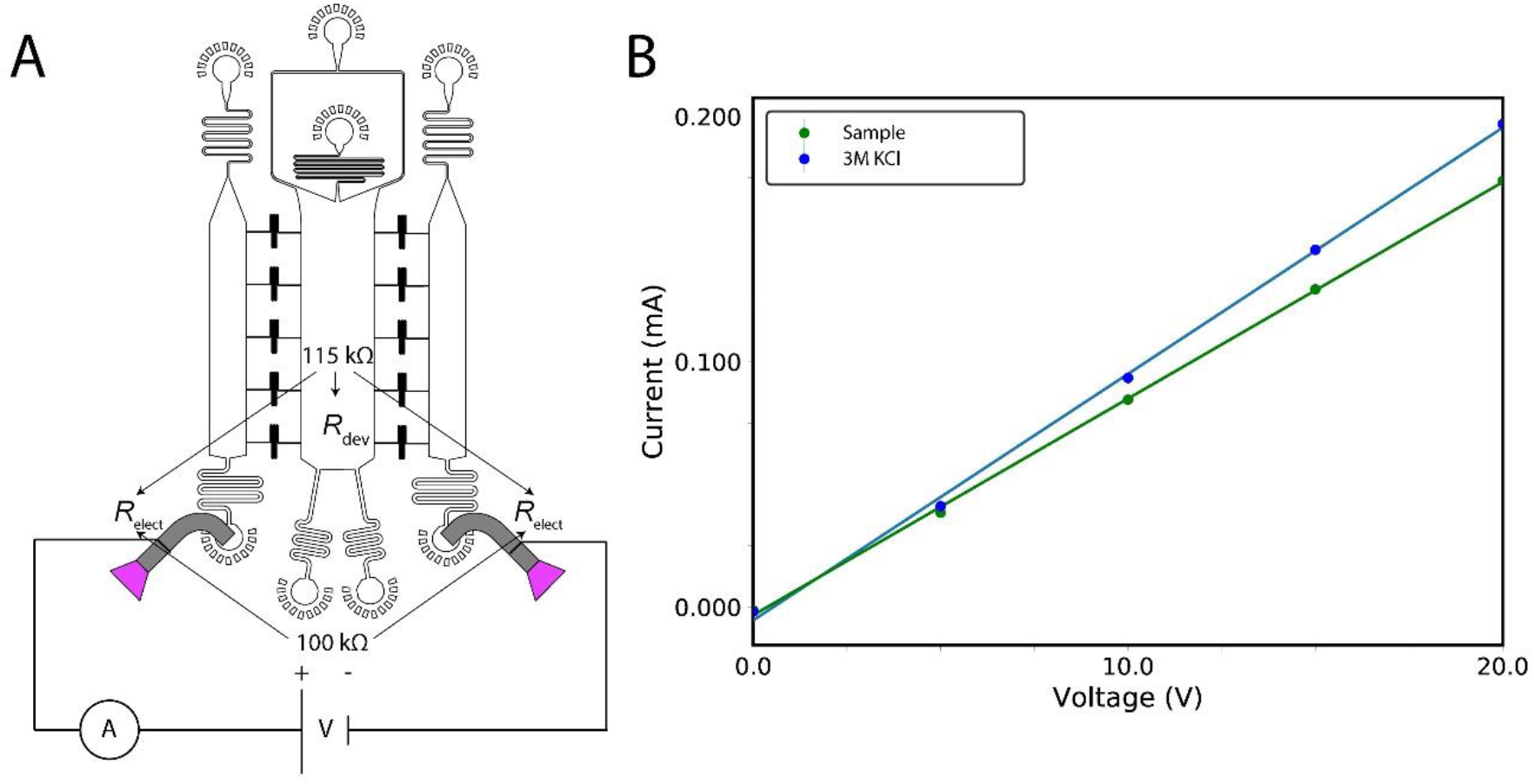
Electrical circuit and calibration of device. **(A)** Circuit schematic displays how the voltage is applied across the μFFE device and indicates the two sources of voltage drop (high electrical resistance), the electrodes (*R*_elect_) and the device itself (*R*_dev_). **(B)** Plot displaying the electrical current transmitted through the device both with the sample present (green) and when the device was filled with 3 M KCl solution (blue). Error bars of three measurements at each voltage are smaller than the marker size. This plot allows for the calibration of the voltage efficiency as described in the text.

### Supplementary Results

**Figure S3.**
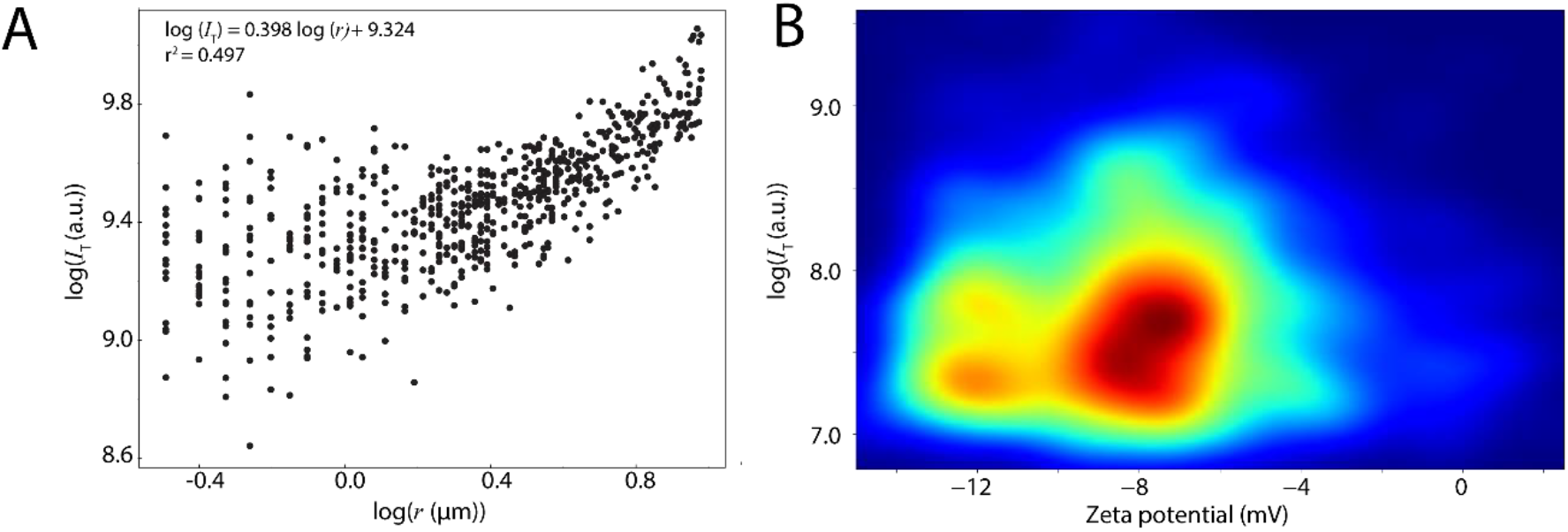
Size dependence of zeta potential. **(A)** Log–log plot of radius, *r*, versus total fluorescence intensity, *I*_T_, of individual FUS condensates at 6 μM FUS in 50 mM TRIS-HCl at pH 7.4 and 50 mM KCl. *r* and *I*_T_ were detected with the TrackMate package in the FIJI image processing software on still images. The correlation between *r* and *I*_T_ of the condensates was fitted with a log model. **(B)**2D plot of zeta potential versus *I*_T_ of individual FUS condensates. The plot shows that condensates with varying zeta potentials have similar distributions of *I*_T_, indicating a lack of correlation between zeta potential and size of condensates.

#### Size dependence of zeta potential

The size of the condensates could not be determined from μFFE experiments because condensates were under flow and appeared blurred in the images due to the 10 ms exposure time. Thus, the size versus zeta potential relationship had to be derived by secondary means. First, static epifluorescence images of FUS condensates were taken. This analysis showed that there is a weak correlation between the total fluorescence intensity (*I*_T_) and the radius (*r*) of FUS condensates (Figure S3A). In a second step, *I*_T_ and zeta potential were derived from images taken during μFFE experiments (Figure S3B). Building on the weak correlation between *I*_T_ and *r*, these data suggest that is no correlation between the zeta potential and *I*_T_, thus indicating that there is no correlation between size and zeta potential.

**Figure S4.**
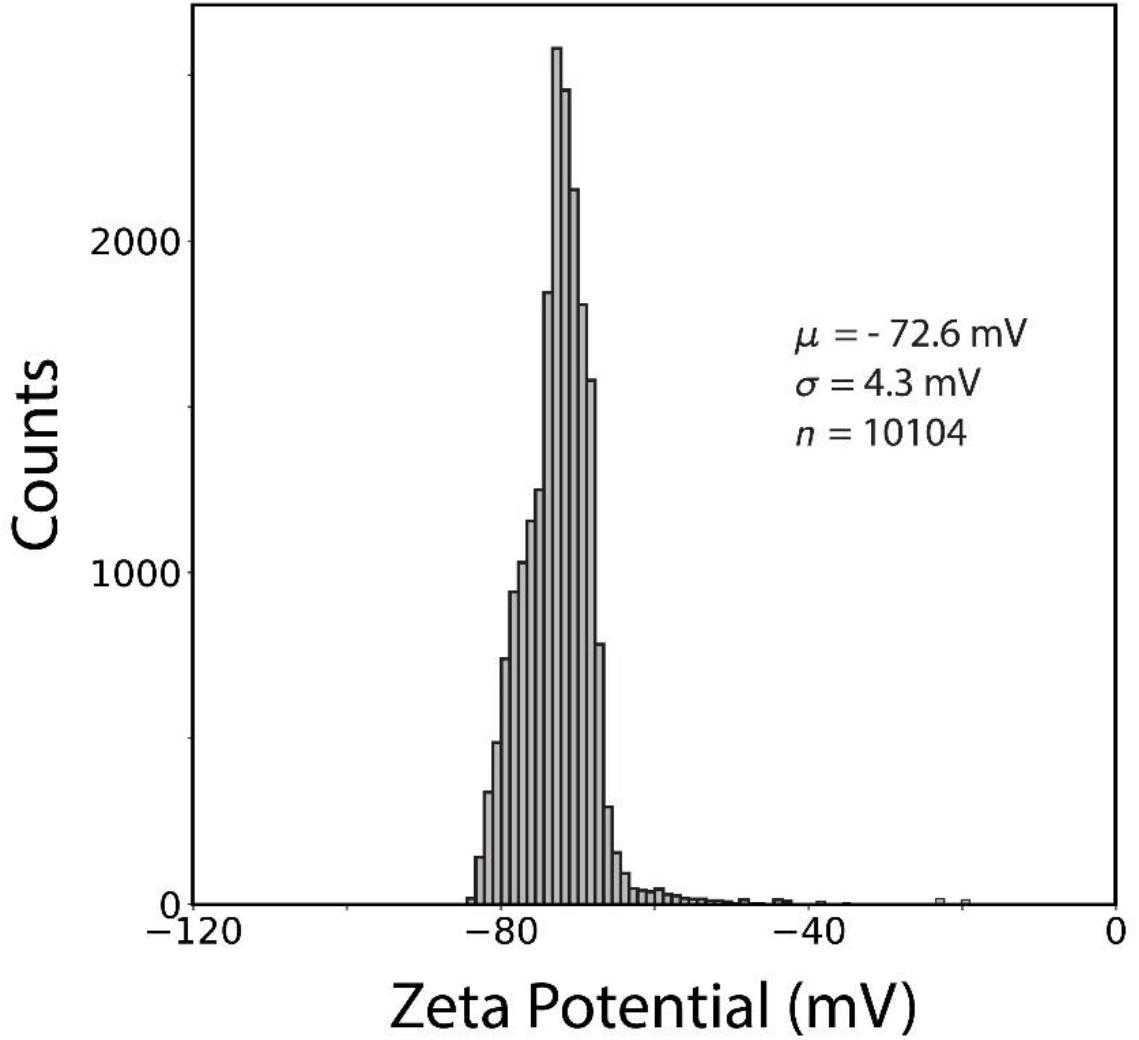
Zeta potential measurement of 60 nm gold nanoparticles. The zeta potential of monodispersed 60 nm gold nanoparticles were measured using the same 3D microfluidic method as for protein condensate measurements.

#### Colloid control measurements

As a control, the zeta potential of 60 nm gold nanoparticles was analyzed using the same microfluidic method as for protein condensate measurements. It was observed that the zeta potential distribution of gold nanoparticles is narrow compared to the distribution of condensates, with the relative standard deviation amounting to only 8% and 20% of that of protein samples. Moreover, there were less broad tails to the distribution in the nanoparticles, which is consistent with the fact that the nanoparticles are monodisperse in size and composition. Note that the nanoparticles are at least an order of magnitude smaller than the condensate systems studied here; hence, due to diffusion effects, the width of the zeta potential distribution of 60 nm gold nanoparticles is likely broader than that of monodisperse particles which are similar in size as condensates.

### Supplementary Computational Methods

#### Coarse-grained protein model

To model the condensation of FUS and PR_25_, we used a reparameterization of the sequence-dependent coarse-grained model of the Mittal group (*12*) that includes enhanced cation–π interactions (*13*). The model treats each amino acid residue as a single bead. Intrinsically disordered regions are modeled as flexible polymers, with inter-residue bonds described using a harmonic potential. Globular regions are treated as rigid bodies. A Coulombic term with Debye–Hückel electrostatic screening was used for long-range electrostatics, while a knowledge-based potential, termed HPS, that is based on a hydrophobicity scale for amino acids developed earlier (*14*) was used to describe pairwise hydrophobic interactions. We have scaled down the set of HPS parameters by 30% to account for the ‘buried’ amino acids contained in the globular “rigid” domains. The model was validated by ensuring that we obtained reasonable qualitative agreement with experiments probing phase behavior of FUS wild type versus FUS prion-like domain (PLD); these experiments reveal a greater propensity for LLPS in the former.

#### Initial atomistic models for coarse-grained simulations

We modelled the full length FUS protein based on Unitprot code K7DPS7 (526 residues, 24 proteins) and a reduced version of the PR_25_ protein (12 Arg and 13 Pro residues alternately positioned, 200 proteins). We developed an atomistic model of FUS by attaching the disordered regions to the resolved structural domains (residues 285–371 (PDB code: 2LCW) and residues 422–453 (PDB code: 6G99)). Initial intrinsically disordered models for PR25 were developed in PyMol (*15*).

#### Minimal coarse-grained model for PolyU

We modelled PolyU (30 strands of 80 nucleotides each) as a flexible polymer that represents each nucleotide as a single bead. Inter-residue bonds were described using a rigid harmonic spring, and long-range electrostatics were modelled using a Coulombic term with Debye–Hückel electrostatic screening plus dispersive interactions. Each bead was assigned a charge of –1 and the HPS set of parameters for Glu dispersive interactions.

#### Coarse-grained simulation methods

We performed direct coexistence simulations at constant volume and temperature to describe the formation of liquid condensates in the different systems. The direct coexistence method consists of simulating both the condensate and diluted phases in the same box separated by an interface. These initial simulation boxes containing both phases were prepared by running simulations at constant temperature and a pressure of 1 bar, using the Berendsen barostat, and then enlarging the simulation box in one direction ~3.5 times. The simulation temperatures were chosen at *T*/*T*_c_ ~0.875, that is T = 350 K for full length FUS and T = 440 K for PR_25_ with polyU. We ran ~2 μs of molecular dynamics simulations using a Langevin thermostat with relaxation time of 5 ps and a time step of 10 fs (*16*). The LAMMPS software molecular dynamics package was used to carry out all the coarse-grained simulations (*17*).

#### Surface tension calculation

We determine the surface tension of both condensates at T/T_c_~0.875, by employing the Kirkwood-Buff expression given in Refs. (*18*, *19*). Our direct coexistence simulations stabilizes two condensate interfaces; thus, the expression for computing the surface tension (*γ*) is: *γ* = *L*_z_ / 2·(*p*_n_–*p*_t_), where *L*_z_ is the length of the box perpendicular to the interface, *p*_n_ is the normal component of the pressure tensor perpendicular to the interface (here *p*_zz_) and *p*_t_ is the average of the tangential components of the pressure tensor (here (*p*_xx_ + *p*_yy_) / 2) (*20*).

#### Back-mapping from coarse grained to atomistic scale

Starting from equilibrium coarse-grained structures of the condensates (*Step 1 of our multiscale procedure*), we built atomic resolution systems following three additional steps. ***Step 2***: We unwrapped the coarse-grained bead coordinates across the periodic boundaries and defined the unwrapped bead positions as coordinates for the amino-acid Cα atoms. Using the tleap module of Amber16 (*21*), we added the missing sidechain and backbone atoms in random orientations. ***Step 3***: Because adding atoms in this way results in significant atomic overlaps that cannot be resolved through standard energy minimization procedures, we mapped these atomistic configurations to the higher-resolution coarse grained model Martini (*22*) and standard Martini Water (*23*). For nucleic acids, the ‘soft’ Martini parameters (*24*) without elastic bonds were used. The system’s energy was then minimized in the Martini resolution for 5000 steps using the steepest descent algorithm. ***Step 4***: Finally, the program “backward” (*25*) was used to backmap the Martini configuration to the atomistic resolution.

#### Atomistic molecular dynamics simulations

After back-mapping, we solvated the atomistic condensates using the Gromacs 2018 command gmx solvate (*26*) with the modified TIP3P water model (*27*) creating a rectangular box with the long side (*z*-direction) 12.5 nm away from the condensate interface. We then added Na^+^/Cl^−^ ions at an initial concentration of 0.2 M using the parameters of Beglov and Roux (*28*) together with the nbfix changes of Luo and Roux (*29*) and Venable et al. (*30*). We used the Charmm36M force field (*31*, *32*), which is one of the standard force field combinations for proteins and nucleic acids in explicit solvent and ions. For the FUS system, this resulted in a system of dimensions 12×12×65 nm with 24 protein molecules (170160 atoms), 250095 water molecules, and 900 Na^+^ and 1236 Cl^−^ ions. For the PR_25_:PolyU system, this resulted in a system of dimensions 7×7×52 nm with 45 protein molecules (21285 atoms), 14 PolyU chains (40 nucleotides, 17906 atoms), 76885 water molecules, and 283 Na^+^ and 277 Cl^−^ ions. All the systems were electro-neutral.

Molecular dynamics simulations were performed with Gromacs 2018 (*26*) using the SETTLE algorithm (*33*) to constrain bond lengths and angles of water molecules and P-LINCS for all other bond lengths, which allowed for a time step of 2 fs to numerically integrate the equations of motions. Temperatures were maintained at 300 K using the v-rescale thermostat (*34*) and the pressure at 1 bar using the Parrinello-Rahman barostat (*35*). Long range electrostatic interactions were calculated using the Particle Mesh Ewald (PME) algorithm (*36*) with a cut-off of 1.0 nm. We first performed a short 25-ns long pre-equilibration molecular dynamics simulation, then after absorption of ions into the condensed phase, the concentration of ions in the diluted phase was verified and adjusted back to 0.2 M NaCl by addition/removal of ions or water molecules. We then conducted a 150 ns long molecular dynamics simulation to investigate the distribution of ions within the condense and diluted phases. The trajectories were analyzed using a combination of Gromacs tools and Python MDAnalysis scripts (*37*). For the calculation of partial densities of atoms across the long box axis, the Cα atoms of the system were first centered within the box and the *density* module of Gromacs was used. For the calculation of interaction preferences, two residues were assumed to be in contact if the minimum distance between their constituent atoms were <3.0 Å in the atomistic resolution and 6.5 Å in the CG resolution. For the calculation of domain interactions, the contacts of all the domain’s constituent residues were summed and normalized by the domain’s length. The trajectories were visualized using VMD (*38*), Pymol (*15*), and Ovito (*39*).

**Figure S5.**
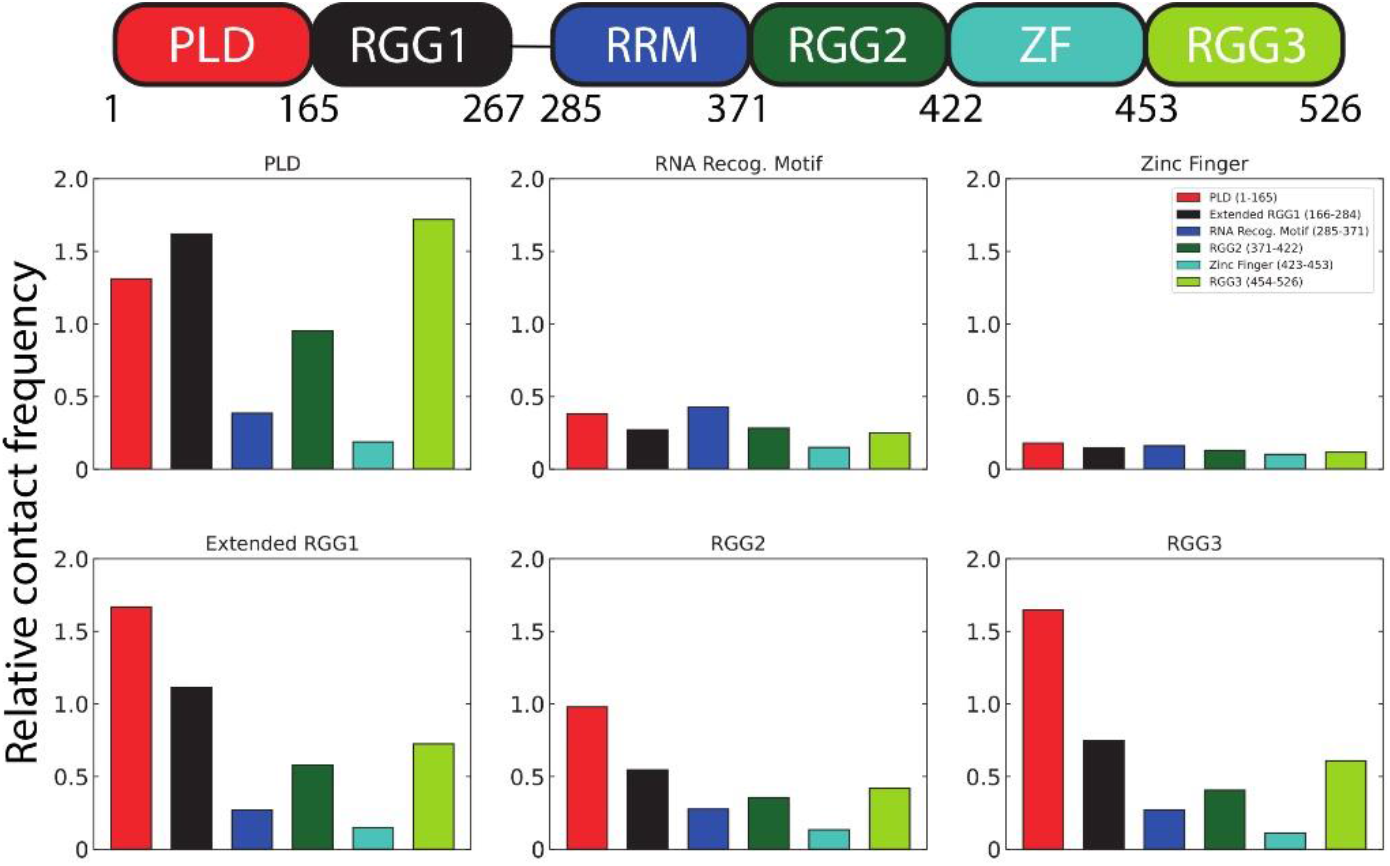
Contacts between FUS domains in condensates. Contact maps showing the relevance of inter-region interactions for each region in FUS (as indicated) within FUS condensates. The bars show the number of inter-protein contacts (amino acids closer than a cut-off of 0.65 nm) mediated by each FUS region normalized by the maximum number of contacts among regions.

**Figure S6.**
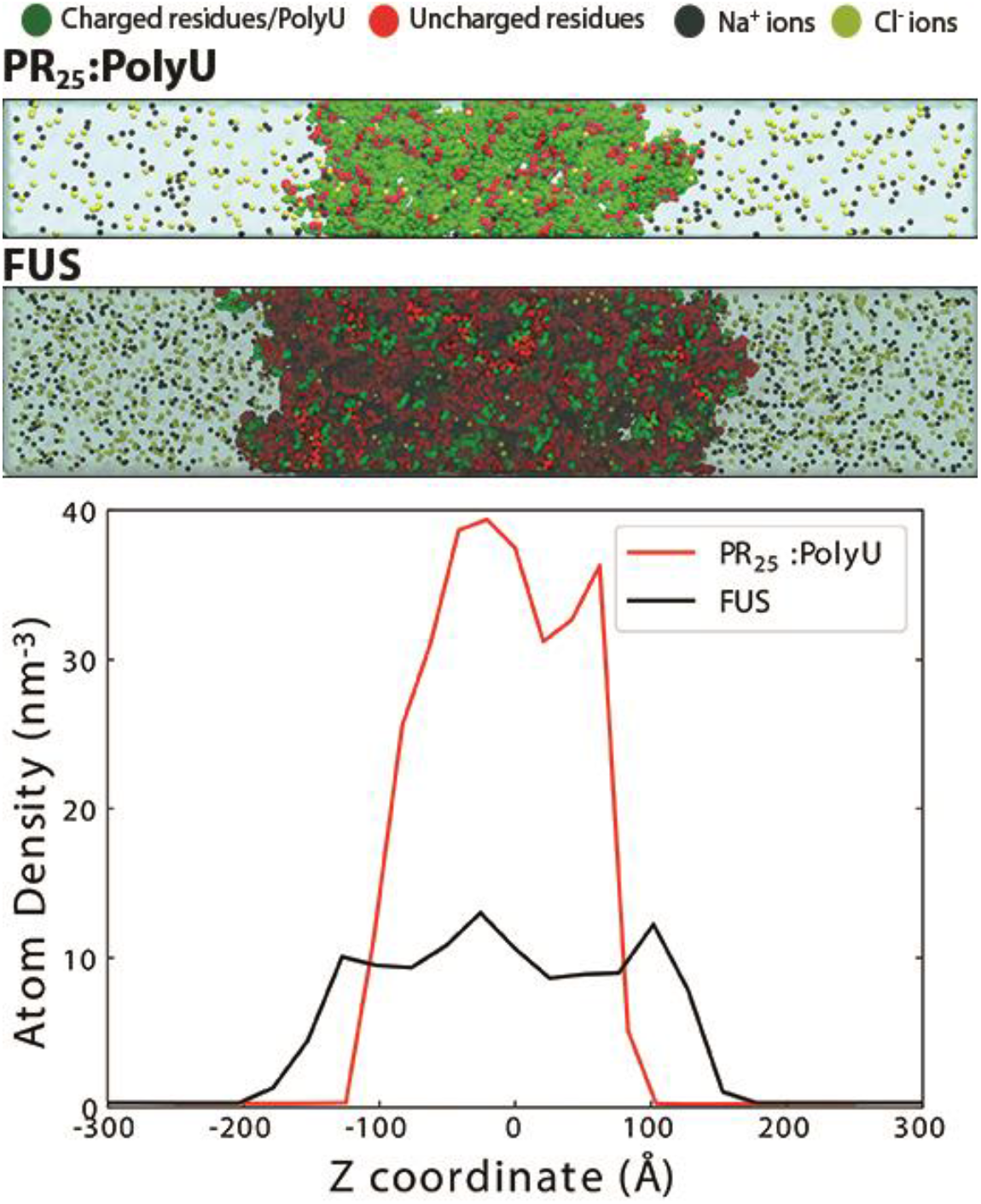
Density of charged atoms in PR25:PolyU and FUS condensates. Total number of charged species (*i.e*., charged atoms and ions) per unit volume (nm^−3^) normalized by the total number of atoms (including ions and water) in each system, as a function of the Z axis. Density profiles were computed from equilibrated atomistic simulations. Snapshots of each system are included in the top panel.

**Figure S7.**
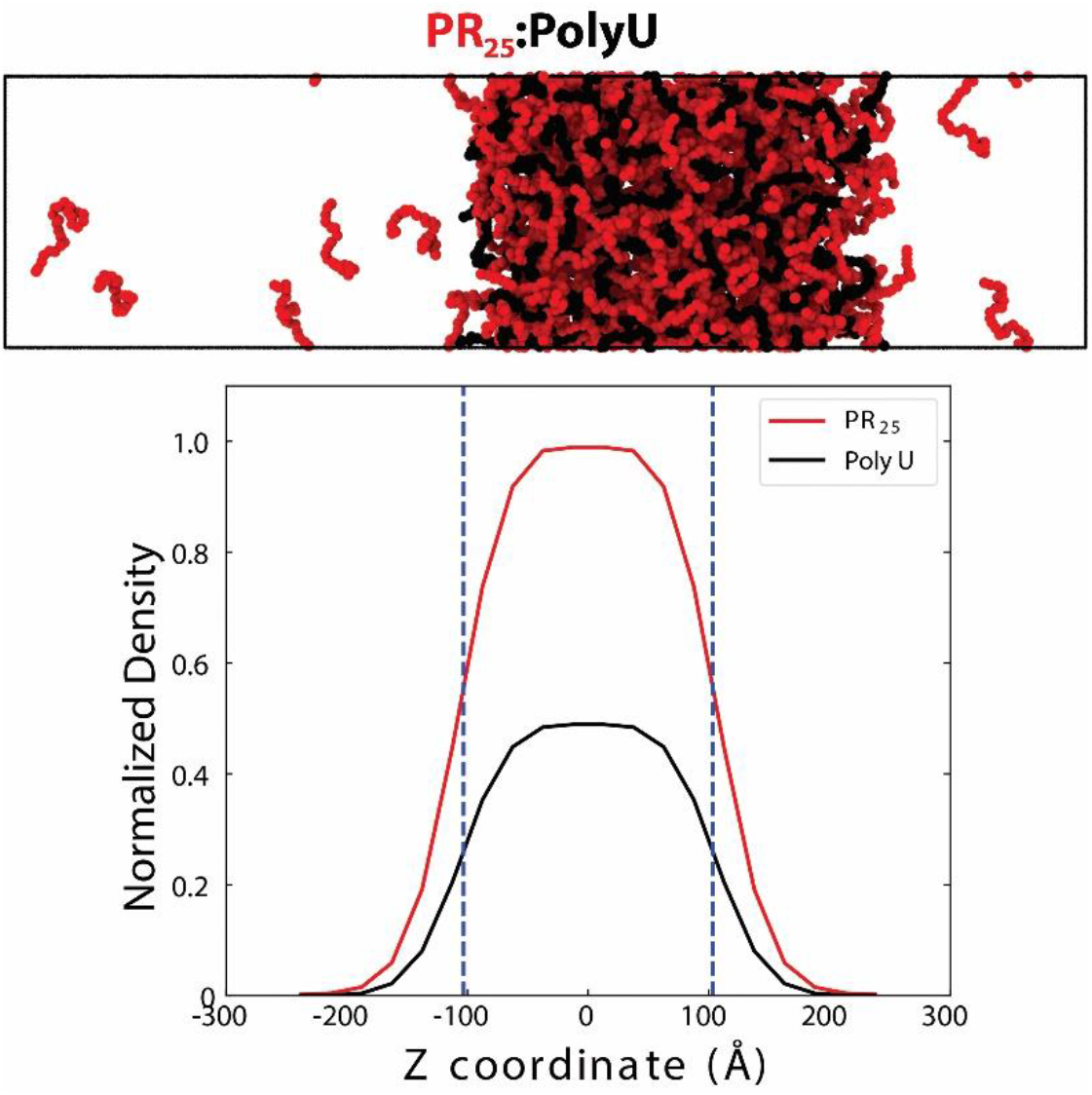
Structure of PR25:PolyU condensates. Normalized densities of individual components (*i.e.*, PR_25_ and PolyU) in PR_25_:PolyU condensates, along the Z coordinate axis. Density profiles suggest that PR_25_ molecules form a monolayer near the condensate interface lowering the condensate surface tension. A snapshot of the system is provided in the top panel.

**Table S1.** Diffusion rates and mobility ratios of ions in condensates. To estimate the differential behavior of ions in and around FUS and PR_25_:PolyU condensates, we measured diffusion coefficients of ions in the condensed (*D*_condensate_) versus the diluted phase (*D*_diluted_) for both systems. The values are calculated from a linear fit of the Mean Square Displacements (MSD) exhibited by the different ions in each phase. The time intervals for the calculation of diffusion coefficients (10 ns) was chosen as the longest interval that minimized intermixing of ions in the condensed phase with ions in the diluted phase. The error estimates are calculated as the difference in diffusion coefficients obtained from two time intervals.

| System | Ions | $D_{\text{condensate}}$<br>(10<sup>-5</sup> cm<sup>2</sup>/s) | $D_{\text{diluted}}$<br>(10<sup>-5</sup> cm<sup>2</sup>/s) | Mobility Ratio<br>($D_{\text{condensate}}/D_{\text{dilut}}$) |
| --- | --- | --- | --- | --- |
| PR<sub>25</sub>:PolyU | Na<sup>+</sup> | 0.866 ± 0.34 | 2.269 ± 0.15 | 0.38 |
|  | Cl<sup>-</sup> | 1.080 ± 0.37 | 2.983 ± 0.66 | 0.36 |
| FUS | Na<sup>+</sup> | 0.511 ± 0.07 | 2.185 ± 0.45 | 0.23 |
|  | Cl<sup>-</sup> | 0.483 ± 0.12 | 3.226 ± 0.24 | 0.15 |

